# Integration of exogenous and endogenous co-stimulatory signals by CAR-Tregs

**DOI:** 10.1101/2022.11.10.516049

**Authors:** Isaac Rosado-Sánchez, Manjurul Haque, Kevin Salim, Madeleine Speck, Vivian Fung, Dominic Boardman, Majid Mojibian, Giorgio Raimondi, Megan K Levings

**Author notes:** **Correspondence:** Megan K. Levings.

## Abstract

Regulatory T cells (Tregs) expressing chimeric antigen receptors (CAR) are a promising tool to promote transplant tolerance. The relationship between CAR structure and Treg function was studied in xenogeneic, immunodeficient mice, revealing advantages of CD28-encoding CARs. However, these models could underrepresent interactions between CAR-Tregs, antigen-presenting cells (APCs) and donor-specific antibodies. We generated mouse Tregs expressing HLA-A2-specific CARs with different costimulatory domains and compared their function in vitro and in vivo. In vitro assays revealed the CD28-encoding CAR had superior antigen-specific suppression, proliferation and cytokine production. In contrast, in vivo protection from skin allograft rejection and alloantibody production was similar between Tregs expressing CARs encoding CD28, ICOS or PD1, but not GITR, 41BB or OX40, co-stimulatory domains. To reconcile in vitro and in vivo data, we analyzed effects of a CAR encoding CD3ζ but no co-stimulatory domain. These data revealed that exogenous co-stimulation via APCs can compensate for the lack of a CAR-encoded CD28 domain. Thus, Tregs expressing a CAR with or without CD28 are functionally equivalent in vivo. This study reveals a new dimension of CAR-Treg biology and has important implications for the design of CARs for clinical use in Tregs.

## INTRODUCTION

Adoptive cell therapy using regulatory T cells (Treg) has emerged as a promising therapeutic strategy to promote transplant tolerance and reduce dependence on immunosuppression (1-3). Multiple clinical studies have demonstrated that polyclonal Treg therapy is feasible, safe and possibly effective (4-7). However, data from pre-clinical models revealed that alloantigen-specific Tregs are significantly more potent at reducing transplant rejection (8, 9). We and others developed a strategy to generate antigen-specific Tregs using Chimeric Antigen Receptors (CARs), synthetic fusion proteins that redirect T cell specificity. CAR-Tregs are more effective than polyclonal Tregs at limiting xenogeneic graft-versus-host disease (xeno-GVHD) (10-12), as well as skin and heart transplant rejection (13-17), and have rapidly transitioned to clinical testing with two ongoing phase I/IIa clinical trials (NCT04817774, NCT05234190) (18).

CARs typically comprise an extracellular single-chain antibody (scFv) domain, a hinge, a transmembrane domain and customizable intracellular signaling domains. They have been extensively studied in the context of cancer immunotherapy, initially as so-called first-generation CARs encoding only a CD3ζ domain, and subsequently as second-or third-generation CARs adding one or more co-stimulatory domains, respectively (19, 20). In the context of transplantation, the optimal CAR design for Tregs is still under debate(8, 21). We recently explored the function of CARs encoding different co-stimulatory domains in human Tregs using an immunodeficient mouse model of xenoGVHD and demonstrated that a second generation CD28-CD3ζ-encoding CAR was optimal in terms of Treg potency, stability and persistence (10). Similar results were found in other studies using PBMC-reconstitution-based, humanized mouse and skin xenograft models (11, 16). However, drawing clinically-relevant conclusions is complicated in these models due to their immunodeficient state and because PBMC reconstitution primarily results in T cell engraftment, with poor/no reconstitution of antigen-presenting cells (APCs), including B cells and dendritic cells (DCs) (22-25).

Suppressing the ability of APCs to activate effector T cells is a primary mechanism by which Tregs maintain peripheral tolerance (2, 26). Tregs suppress APCs using a range of strategies including CTLA-4-mediated transendocytosis of CD80/86 (27, 28), trogocytosis of MHC class II (29, 30) and suppression of cytokine production (31). Tregs also control the generation of donor-specific anti-HLA antibodies (DSA) by directly suppressing B cell function (32, 33), inducing B cell apoptosis (32) and/or inhibiting follicular helper cells (34-38). APCs (39-41) and DSAs (42, 43) both have critical roles in transplant rejection, so identifying the optimal CAR-Treg design to regulate these cells and processes is an important outstanding question (44).

In this study, we used a mouse model of HLA-A2^+^ skin transplantation to study the structure-function relationship of CAR-Tregs. HLA-A2-specific CARs with different co-stimulatory domains were expressed in Tregs and studied in vitro and in vivo in an immunocompetent setting. We explored how CAR-Tregs integrate signals from exogenous and endogenous sources and how signal origin shapes function.

## RESULTS

### Generation of co-stimulation domain variant CAR-Tregs in mice

We generated eight HLA-A2-specific CAR variants containing different transmembrane and co-stimulatory domains derived from CD28 and TNFR family proteins that have relevance to Treg biology (45) (**Fig 1A**). CAR variants were cloned into a bicistronic retroviral vector upstream of a mKO2 reporter. CD4^+^CD8^−^Thy1.1^+^Foxp3^gfp+^ Tregs were sorted, polyclonally stimulated, transduced and expanded (**Supplemental Fig 1**). Control Tregs and conventional T cells (Tconvs) were expanded in a similar manner but transduced with an irrelevant CAR or left untransduced.

**Figure 1.**
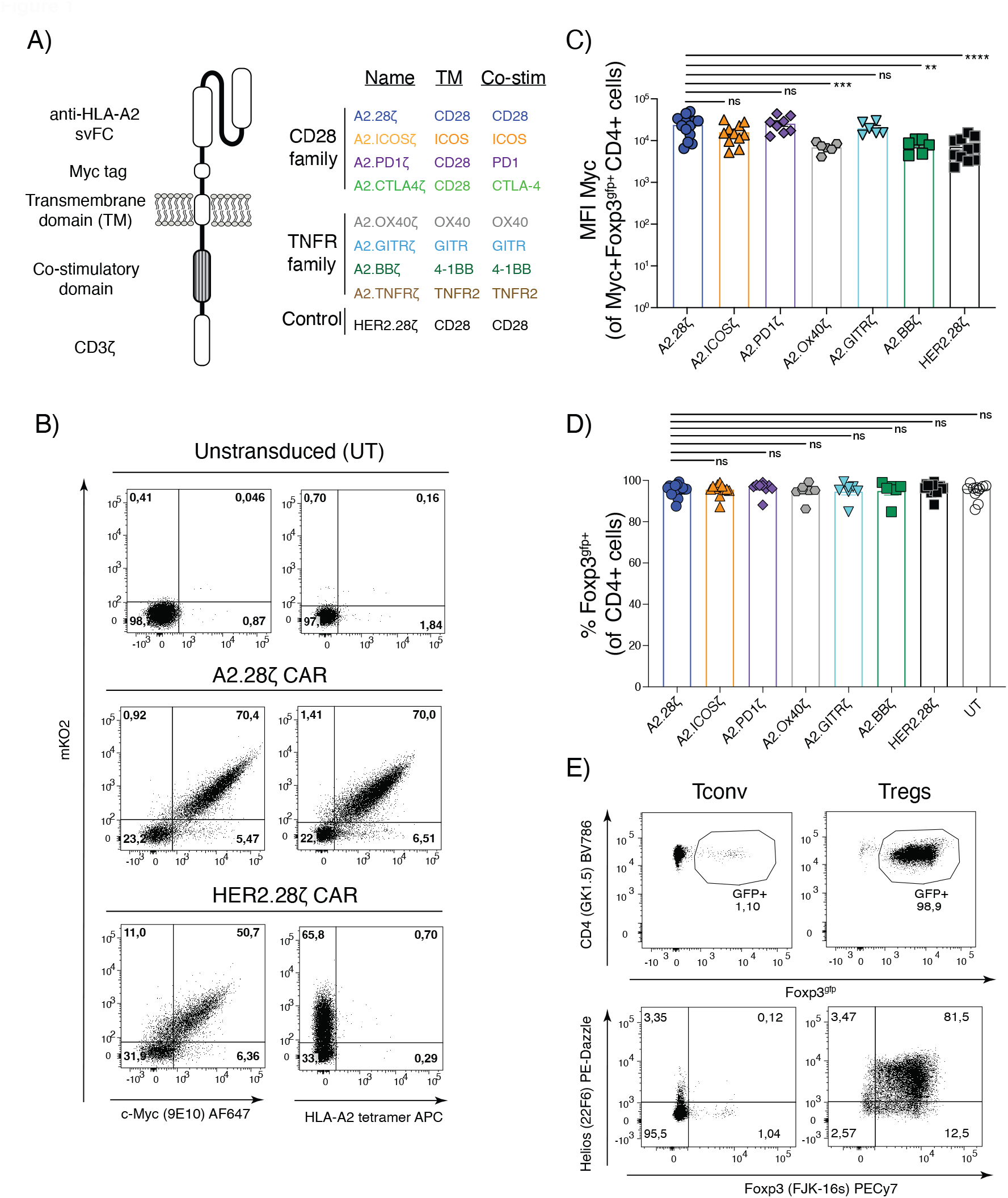
Design and expression of co-stimulatory domain CAR variants. (**A**) Schematic diagram summarizing the transmembrane and signaling domains incorporated into the second-generation CAR variants. (**B**) Representative flow cytometry plots showing CAR (c-Myc) and mKO2 reporter expression, and binding to an HLA-A2 tetramer. (**C**) MFI of CAR expression for different CAR variants in Tregs after expansion gated on live Myc^+^CD4^+^Foxp3^gfp+^ cells. (**D**) Foxp3^gfp^ expression in Tregs after expansion, gated on live CD4^+^ cells. (**E**) Representative data for intracellular Foxp3 and Helios expression in CAR-Tregs and control Tconvs after expansion, gated on total live CD4^+^ cells. Data pooled from at least 6 independent experiments and shown as mean±SEM. Statistical significance was determined using one-way ANOVA with a Holm-Sidak post-test, *p<0.05, **p<0.01, ***p<0.001, ****p<0.0001.

Treg transduction and CAR expression were measured by expression of the CAR-encoded extracellular Myc tag and mKO2 (**Fig 1B; Supplemental Fig 2A**). With the exception of CTLA4- and TNFR2-encoding CARs (**Supplemental Fig 2B**, not analyzed further), CAR variants were detected on the cell surface and bound to HLA-A2 tetramers (**Fig 1B**). After expansion, on average, ∼70% of cells expressed a CAR (**Supplemental Fig 2C**). Expression levels of OX40- and 41BB-encoding CARs, and the control HER2 CAR, were lower than the CD28-encoding CAR (**Fig 1C**), but there were no differences in gfp (Foxp3 reporter) or intracellular Foxp3 expression, demonstrating high Treg purity following transduction and expansion (**Fig 1D&E**).

### Co-stimulatory CAR variants differ in their ability to stimulate Tregs

To assess CAR variant function, CAR-Tregs were labelled with CPDeF450 and co-cultured with irradiated HLA-A2^pos^ K562 cells for 72 hrs. Only Tregs expressing a CAR proliferated in response to HLA-A2 (**Fig 2A**). Differences in Treg proliferation were observed, with the CD28-encoding CAR inducing the strongest proliferative response, followed by the ICOS-encoding CAR (**Fig 2B**). TNFR-family co-stimulatory CARs (OX40, GITR and 4-1BB) induced a moderate proliferative response (**Fig 2B**), whilst the PD1-encoding CAR induced little proliferation, corroborating our previous study in human Tregs (10) and other studies in CAR-T cells (46).

**Figure 2.**
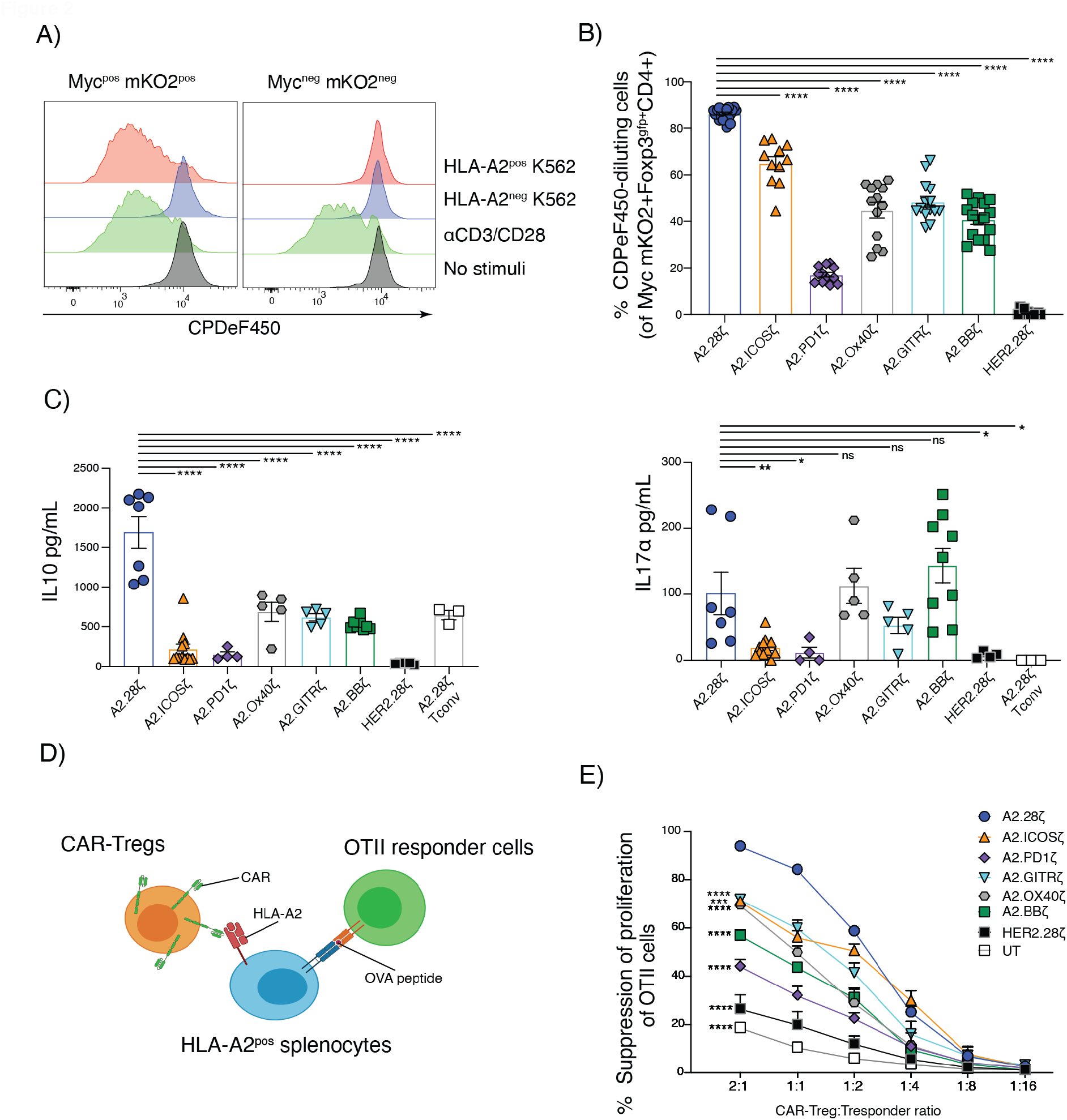
Co-stimulatory CAR variants differ in their ability to stimulate Tregs. (**A**,**B&C**) Tregs expressing the indicated CAR were stained with CPDe450 and co-cultured with HLA-A2^pos^ or HLA-A2^neg^ K562 cells, polyclonal stimulated with anti-CD3/28, or left unstimulated for 3 days. (**A**) Representative histograms from A2.28ζ-CAR Tregs comparing proliferation of gated CAR+ (Myc^+^mKO2^+^) or CAR-negative (Myc^-^mKO2^-^) cells. (**B**) Frequencies of CAR-Tregs that divided following 3-day co-culture with HLA-A2^+^ K562s, determined by CPDeF450 dilution, gated on cMyc^+^mKO2^+^Foxp3^gfp+^CD4^+^ cells. (**C**) Cytokine secretion following 3-days of co-culture with HLA-A2^pos^ K562s. (**D&E**) CAR-Tregs were co-cultured with OTII CD4^+^ T cells at varying ratios in the presence of irradiated HLA-A2^+^ splenocytes and OVA peptide. (**D**) Schematic diagram of linked suppression assay. (**E**) CAR-Treg mediated suppression of the OTII CD4^+^ T cell proliferation, as determined by Ki67 expression. UT = Untransduced. Data pooled from 5 (**B**), 3 (**C**) and 2 (**E**) independent experiments, shown as mean±SEM. Statistical significance was determined using one-way (**B/C**) or two-way (**E**) ANOVA with a Holm-Sidak post-test comparing to CD28-based CAR-Tregs, *p<0.05, **p<0.01, ***p<0.001, ****p<0.0001.

Analysis of cell culture supernatants revealed that Tregs expressing the CD28-encoding CAR secreted the highest levels of IL10. CARs encoding TNFR family domains (OX40, GITR, 4-1BB) induced medium levels of IL10 and PD1- and ICOS-encoding CARs induced the lowest (**Fig 2C-left**). Low levels of IL17A were secreted by Tregs expressing the ICOS- and PD1-encoding CARs, contrasting with a previous study performed with CAR-T cells that showed an ICOS-encoding CAR induced high IL17A production (47) (**Fig 2C-right**). In comparison to A2-CAR T cells, none of the CAR-Tregs variants secreted significant amounts of pro-inflammatory cytokines or IL-2 (**Supplemental Fig 3A**).

To test how CAR signaling influenced Treg function, antigen-dependent, linked suppression assays were performed where the ability of CAR-Tregs to inhibit OTII CD4^+^ T cell proliferation was measured (**Fig 2D**). Tregs expressing the CD28-based CAR exhibited the greatest suppressive function (**Fig 2E; Supplemental Fig 3B**). Tregs expressing the other CARs varied in their suppressive capacity. PD1-encoding CAR-Tregs were the least suppressive, but remained more suppressive than the polyclonal HER-CAR or untransduced Treg controls. Thus, as we previously found in human CAR-Tregs(10), in an in vitro setting, CAR-Treg activation is strongly influenced by the CAR co-stimulatory domain.

### In vivo effects of Tregs expressing co-stimulatory CAR variants on skin rejection

We next compared the function of CAR-Treg variants using an immunocompetent mouse model of allogeneic skin transplantation (15). Wild-type Bl/6 mice received HLA-A2^+^ Bl/6 skin grafts and were intravenously administered with 1×10^6^ CAR-Tregs. Consistent with our previous study (15), CAR-Tregs delayed, but did not prevent skin rejection: median survival time was 20 days for mice treated with A2.CD28ζ CAR-Tregs versus 14 days for PBS (**Fig 3A-left**). CAR-Tregs encoding other CD28-family domains, ICOS or PD1, also delayed skin rejection with median survival times of 20 days for ICOS and 19.5 for PD1. On the other hand, with the exception of GITR, Tregs encoding CARs with TNFR family-derived domains failed to extend graft survival. The median survival times were 14 days for OX40-, 17 days for 41BB- and 19.5 days for GITR-encoding CAR-Tregs (**Fig 3A-right**). Overall, no other CAR-Treg extended graft survival significantly longer than A2.CD28ζ CAR-Tregs (**Supplemental Table 1**).

**Figure 3.**
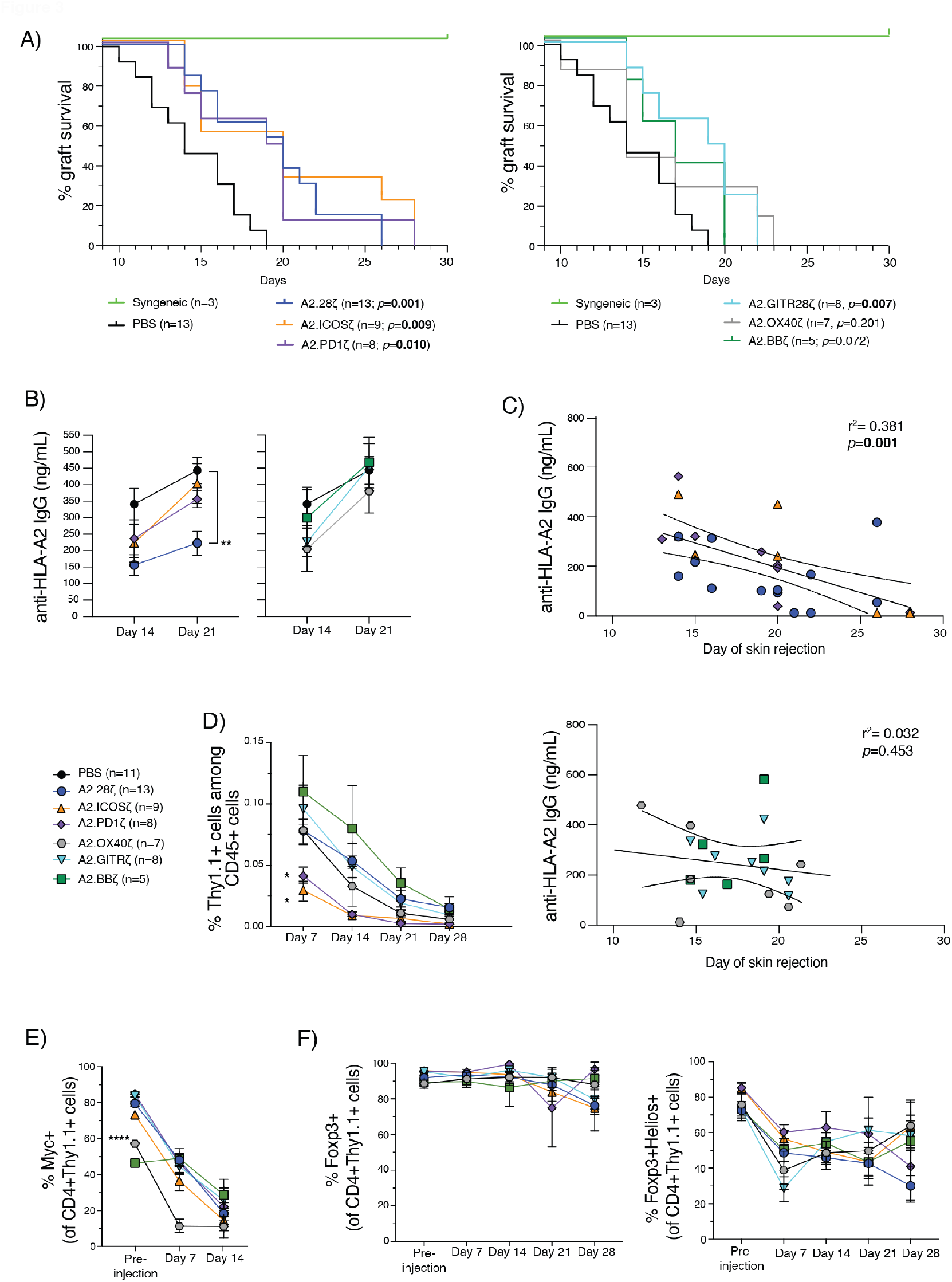
In vivo effects of Tregs expressing co-stimulatory CAR variants on skin rejection. BL/6 mice were transplanted with skin grafts from syngeneic or HLA-A2^+^ BL/6 mice and intravenously administered 1×10^6^ CAR-Tregs. (**A**) Skin graft survival curves and (**B**) levels of anti-HLA-A2 IgG Abs from mice infused with Tregs expressing CARs encoding costimulatory domains from CD28 (left) or TNFR (right) family members. For A, data from mice receiving no Treg treatment (PBS) or transplanted with syngeneic wild-type BL/6 grafts are shown in both graphs. (**C**) Correlation between anti-HLA-A2 IgG antibodies in plasma at day 14 and skin graft rejection of mice receiving Tregs bearing CD28 and TNFR family-based CARs. (**D**) % Thy1.1^+^ CAR-Tregs of total CD45^+^ T cells in peripheral blood over time. (**E-G**) Phenotype of Thy1.1^+^CD4^+^ CAR Tregs in peripheral blood over time including expression of: (**E**) CAR (Myc+) (**F**), FoxP3 alone (left) and FoxP3 with Helios (right). Data are mean±SEM pooled from 4-individual experiments. Statistical significance was determined using log-rank Mantel-Cox test (**A**), two-way ANOVA (**B**,**D-F**) with a Holm-Sidak post-test, and Pearson correlation (**C**). *p<0.05, **p<0.01, ***p<0.001, ****p<0.0001.

DSAs are important mediators of organ rejection (42) so CAR-Treg control of anti-HLA-A2 IgG generation was assessed. Mice treated with A2.CD28ζ CAR-Tregs had significantly lower levels of anti-HLA-A2 IgG compared to PBS mice (**Fig 3B**), corroborating our previous observations (15). Conversely, no other CAR-Treg tested significantly reduced the levels of anti-HLA-A2 IgG compared to PBS mice. Seeking to assess if there was a correlation between control of anti-HLA-A2 IgG and graft rejection, a regression analysis was performed revealing a negative correlation between amounts of anti-HLA-A2 IgG and graft survival (**Supplemental Fig 4A**). Interestingly, when this analysis was performed separately for CD28-versus TNFR-family encoding CARs, the correlation was only present for the former (**Fig 3C**).

CAR-Treg persistence and phenotype were tracked in blood on a weekly basis. Only Tregs expressing ICOS-or PD1-encoding CARs persisted significantly less than A2.CD28ζ CAR-Tregs (**Fig 3D; Supplemental Fig 4B**). With the exception of the A2.OX40ζ CAR, there were no differences in the proportion of CAR-expressing Tregs (**Fig 3E**). Expression of Foxp3 and Helios were equivalent between all CAR-Treg groups, showing that none of the co-stimulatory domains negatively affected Treg stability *in vivo* (**Fig 3F**).

### A co-stimulatory domain is dispensable for CAR-Treg function in vivo

The minimal differences between some CAR-Tregs variants in this immunocompetent setting contrast with the in vitro data in this study and previous studies that used immunodeficient mouse models, both of which clearly showed a superior function of CD28-encoding CARs (10, 11, 16, 48). Seeking to understand the mechanistic basis for these findings, we hypothesized that CAR-Tregs may receive additional signals in vivo that compensated for weaker CAR-mediated activation. To address this possibility, we tested a first-generation CAR (A2.ζ CAR) that lacked a co-stimulatory domain (**Fig 4A**) and performed direct comparisons with the second-generation A2.28ζ CAR. The A2.ζ and A2.28ζ CARs were expressed at similar levels and no differences in Foxp3^gfp^ expression were observed (**Supplemental Fig 5A**). In vitro assays revealed that upon stimulation with HLA-A2^pos^ K562 cells, A2.ζ CAR-Tregs had significantly lower proliferation and cytokine secretion than A2.28ζ CAR-Tregs (**Fig 4B; Supplemental Fig 5B**). When tested in the OTII linked suppression assay there was a trend to lower antigen-specific suppression with A2.ζ compared to A2.28ζ CAR-Tregs (**Fig 4C; Supplemental Fig 5C**).

Following adoptive transfer into an immunocompetent skin transplant model, A2.ζ and A2.28ζ-CAR Tregs were equal in their ability to delay skin rejection (median survival time of 20 days for both vs 14 days for PBS mice) (**Fig 4D**). Additionally, A2.ζ and A2.28ζ CAR Tregs were similarly able to suppress the generation of anti-HLA-A2 IgG antibodies (**Figure 4E**) and there was a strong correlation between anti-HLA-A2 IgG levels and graft rejection (**Supplemental Fig 6A**). There was a trend, although not significant, to lower persistence of A2.ζ CAR-Tregs (**Fig 4F; Supplemental Fig 6B**) and no differences in expression of the CAR (**Fig 4G**), Foxp3 and Helios (**Fig 4I**) or functional markers (**Supplemental Fig 6C**). Together, these results suggest CAR-encoded co-stimulation is redundant for CAR-Treg function in an immunocompetent model.

**Figure 4.**
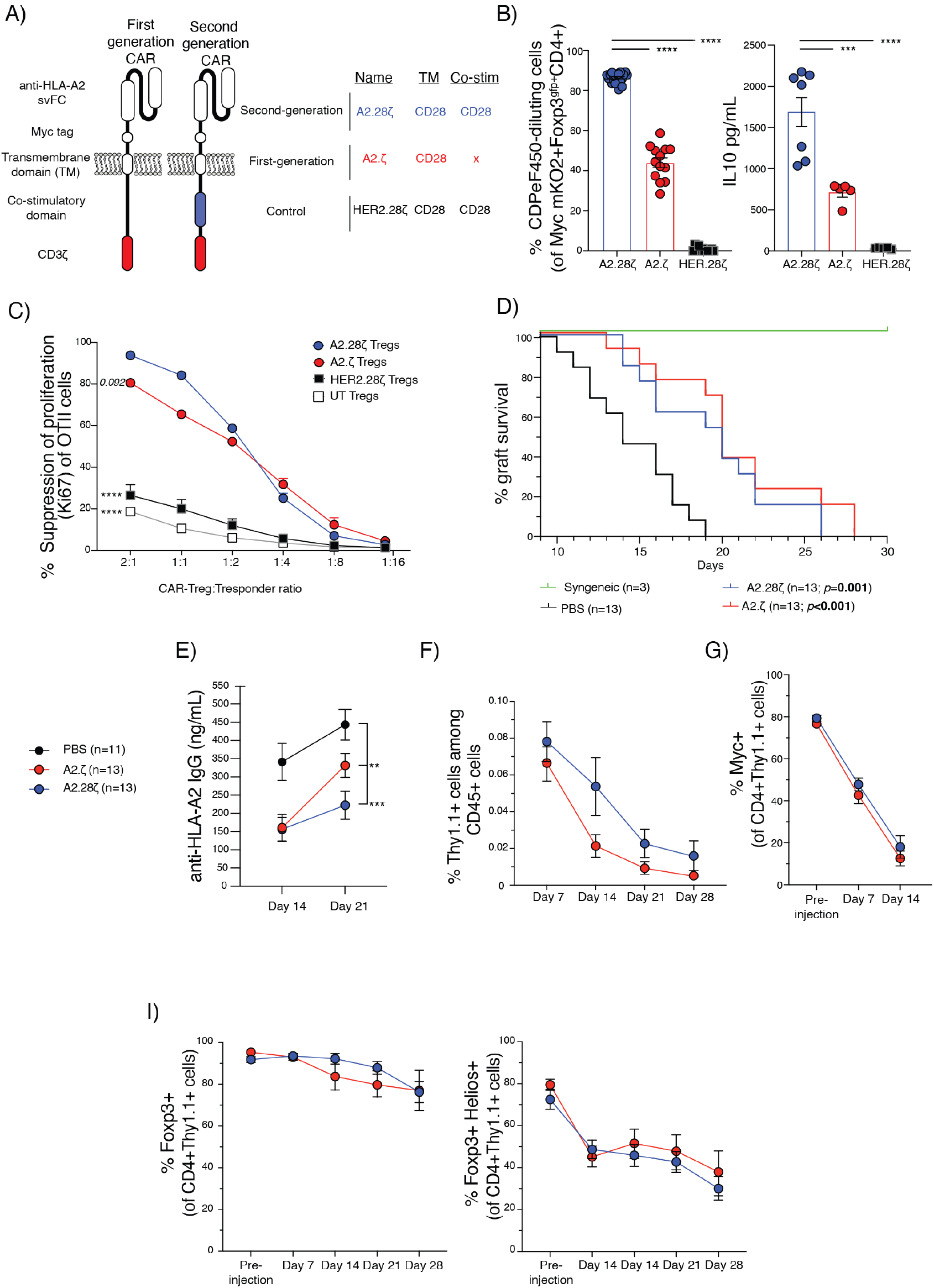
A CAR co-stimulatory domain is dispensable for CAR-Treg in vivo. (**A**) Schematic diagram of the first- and second CAR used. Tregs expressing the indicated CARs were stained with CPDe450 and co-cultured with HLA-A2^pos^ for 3 days. (**B**) % CAR-Tregs that divided, determined by CPDeF450 dilution (left) and IL10 secretion (right), measured in culture supernatants. (**C**) CAR-Tregs were co-cultured with OTII CD4+ T cells at varying ratios in the presence of irradiated HLA-A2^+^ splenocytes and OVA peptide. CAR-Treg mediated suppression of the OTII CD4^+^ T cell proliferation, as determined by Ki67 expression. UT = Untransduced. (**D-I**) BL/6 mice were transplanted with skin grafts from syngeneic or HLA-A2^+^ BL/6 mice and intravenously administered 1×10^6^ CAR-Tregs. (**D**) Skin graft survival curves and (**E**) levels of anti-HLA-A2 IgG Abs from mice infused with Tregs expressing first- and second-generation CARs. (**F**) % Thy1.1^+^ CAR-Tregs of total CD45^+^ T cells in peripheral blood over time. (**G-I**) Phenotype of Thy1.1^+^CD4^+^ CAR Tregs in peripheral blood over time including expression of: (**G**) CAR (Myc+) (**I**), FoxP3 and FoxP3 and Helios. Data are mean±SEM pooled from 2 (**C**), 3 (**B**, right & **D**-**I**) or 5 (**B**, left) individual experiments. Data from the A2.28ζ, HER2.28ζ and UT conditions are also shown from Figures 1C&E and 3A,B,D, E&F. Statistical significance was determined using one-way (**B**) or two-way (**C, E-I**) ANOVA with a Holm-Sidak post-test or log-rank Mantel-Cox test (**D**), *p<0.05, **p<0.01, ***p<0.001. ****p<0.0001.

### CAR-Tregs integrate exogenous and CAR-encoded co-stimulation

A fundamental difference between our in vivo studies and those previously performed with humanized mice is that the latter lacks professional APCs. As such, we hypothesized that in an immunocompetent in vivo setting, CD28 naturally-expressed by the CAR-Tregs may engage CD80/86 on APCs and compensate for a weak/absent CAR-encoded co-stimulatory signal. To investigate this possibility, A2.ζ-or A2.28ζ-CAR Tregs were stimulated with HLA-A2^pos^CD86^neg^ or HLA-A2^pos^CD86^pos^ K562s, after which proliferation and activation were determined by the expression of Ki67, CTLA-4, PD1 and LAP (**Fig 5A**). In the absence of exogenous CD86, A2.28ζ-CAR Tregs were significantly more activated and proliferative than A2.ζ-CAR Tregs (**Fig 5B; Supplemental Fig 7A**). However, in presence of CD86 these differences diminished with the first- and second-generation CAR-Tregs showing similar levels of activation and proliferation (**Fig 5B; Supplemental Fig 7A**). To further validate these findings, Tregs were stimulated with HLA-A2^pos^CD86^pos^ K562s in the presence of CTLA4-Ig to block CD86. CTLA4-Ig treatment reduced the proliferation and activation of A2.ζ-CAR Tregs to similar levels found in absence of CD86 (**Fig 5B&C; Supplemental Fig 7A&B**). The inhibitory effect CTLA-Ig was overcome by adding an agonistic anti-CD28 mAb able to induce CD28-signaling independently of CD86.

**Figure 5.**
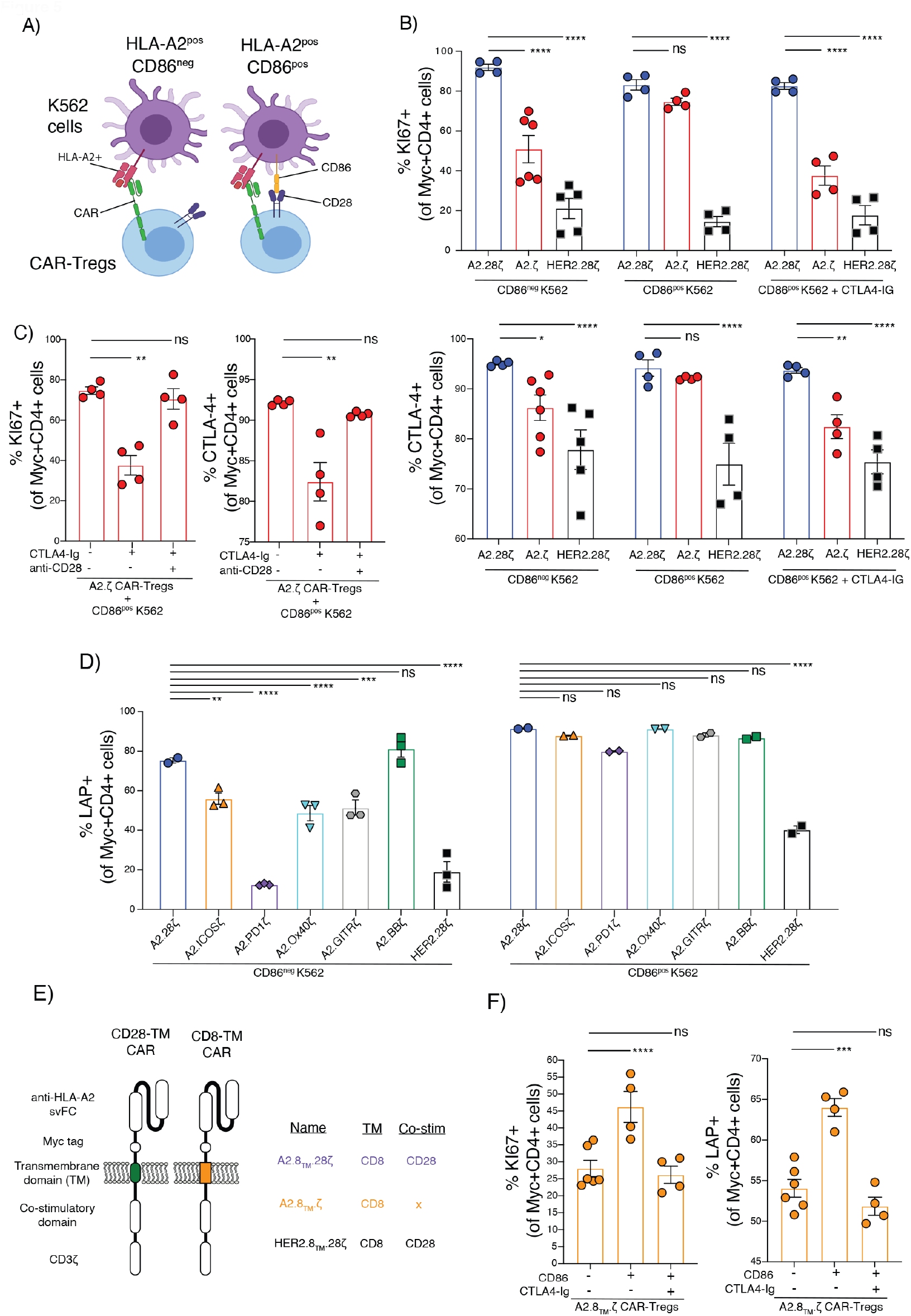
Effect of exogenous co-stimulation on CAR-Tregs. CAR-Tregs were co-cultured with CD86^pos^HLA-A2^pos^ or CD86^neg^HLA-A2^pos^ K562 cells at a 1:2 K562:Tregs ratio for 3 days. (**A**) Schematic diagram of assay. (**B-C**) Ki67 and CTLA4 expression in CAR-Tregs following 3-days of co-culture, gated on Myc+CD4+ live cells. (**D**) LAP expression in different co-stimulatory-encoding CAR-Tregs following 3-days of co-culture, gated on Myc+CD4+ live cells. (**E**) Schematic diagram of the CD8α-transmembrane domain (TM) CARs generated. (**F**) Ki67 and LAP expression in first-generation CD8α-TM CAR-Tregs following 3-days of co-culture, gated on Myc+CD4+ live cells. Where indicated, CTLA4-Ig and an anti-CD28 agonist mAb were added at 10 μg/mL. Data are mean±SEM pooled from 2 (**B**,**C**) and 1 (**D**,**F**) independent experiments. Statistical significance was determined using one-way (**C/F**) or two-way (**B**,**D**) ANOVA with a Holm-Sidak post-test, *p<0.05, **p<0.01, ***p<0.001, ****p<0.0001..

Having shown that co-stimulation through native CD28 can act in conjunction with CAR-mediated CD3ζ signaling to fully activate Tregs, we next asked how CD28 signaling combines with signals from other co-stimulatory domain CARs. Corroborating our previous findings, Tregs encoding different CD28-or TNFR-family CARs were activated to differing degrees upon co-culture with HLA-A2^pos^CD86^neg^ K562s. However, these differences were reduced in the presence of HLA-A2^pos^CD86^pos^ K562s, demonstrating that the function of CAR-Tregs is influenced by both CAR stimulation and CD86 engagement (**Fig 5D**; **Supplementary Fig 8A**).

It has previously been shown that CARs encoding a CD28-derived transmembrane (TM) domain can dimerize with endogenous CD28(49). To exclude the possibility that our observations could be related to interactions between the CAR CD28 TM and native CD28, resulting in the presence of a CD28 signal even in the absence of a CAR-encoded CD28 endodomain, we generated new CARs encoding a CD8α-derived TM (**Fig 5E**). Tregs expressing the indicated CARs were stimulated with HLA-A2^pos^CD86^neg^ or HLA-A2^pos^CD86^pos^ K562 cells, revealing that first-generation CARs with CD8α TM domains are similarly able to respond to exogenous CD28 stimulation (**Fig 5F**; **Supplemental Fig 8B**) suggesting a minimal impact of the type of TM in this process.

To further confirm that co-stimulatory domains in a CAR are dispensable for Tregs if co-stimulation is provided by natural APCs (i.e. rather than K562 cells), we analyzed the ability of A2.28ζ and A2.ζ CAR-Tregs to inhibit the antigen-presenting capacity of DCs. CAR-Tregs were co-cultured with HLA-A2^+^CD11c^+^ DCs for 24-48 hrs, after which the expression of CD80, CD86 and MHC class-II in the DCs was assessed (**Fig 6A-B**). A2.28ζ- and A2.ζ-CAR Tregs were equally able to suppress CD80, CD86 (**Fig 6C**) and MHC-II expression (**Supplementary Fig 9A**). This effect was consistent at different time points and different CAR-Tregs/DCs ratios (**Supplemental Fig 9B**). In concordance with previous results, the suppressive effect of A2.ζ-CAR Tregs was strongly inhibited by CTLA4-Ig (**Fig 6D**).

**Figure 6.**
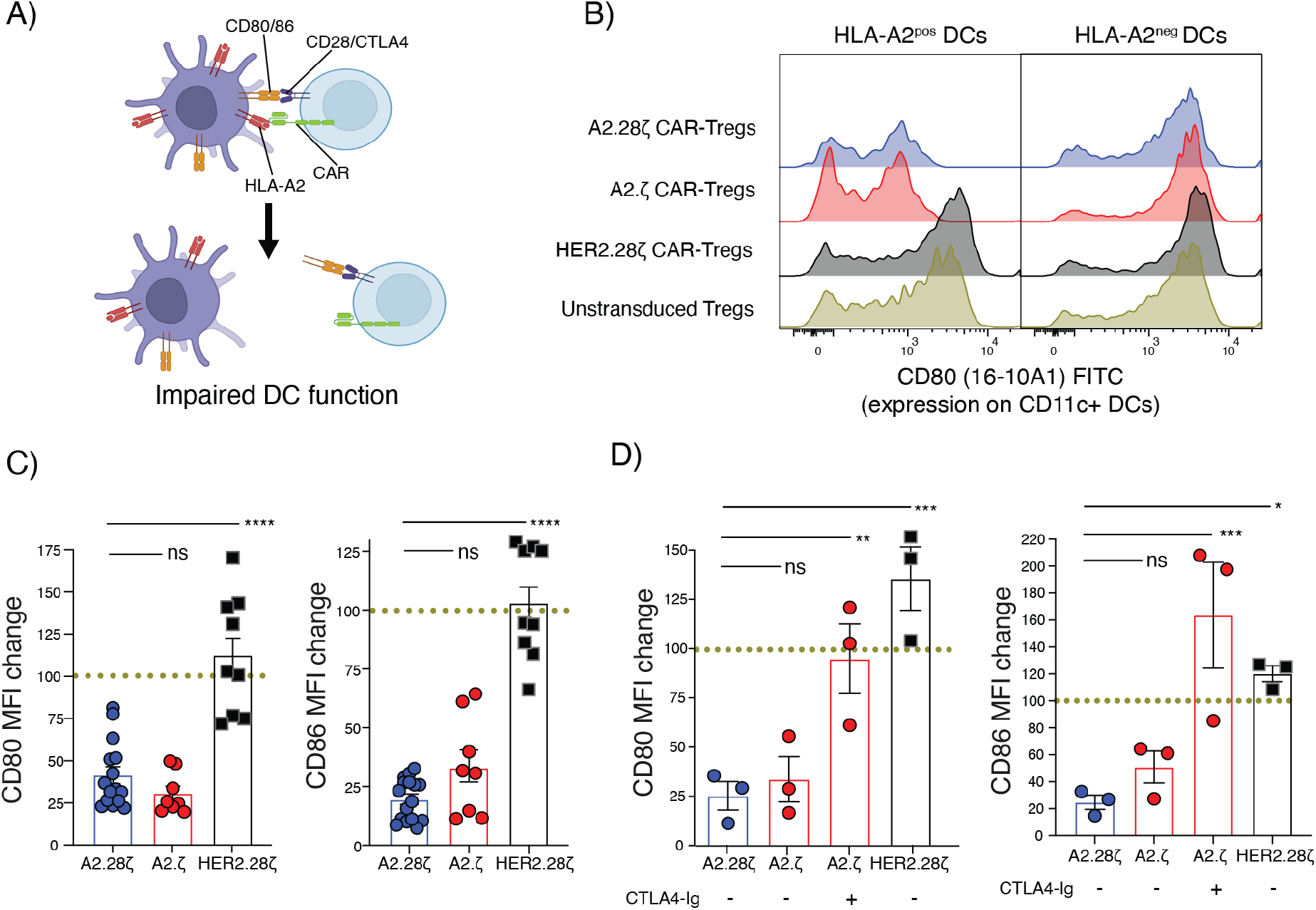
APC suppression by first- and second-generation CAR-Tregs. CAR-Tregs were co-cultured with splenic HLA-A2^+^ CD11c^+^ DCs at a 1:2 or 1:5 DCs:Tregs for 1 or 2 days. (**A**) Schematic of DC suppression assay. (**B**) Representative histograms showing CD80 expression on CD11c^+^ DCs after 2-days of cultured with the indicated types of Tregs. (**C**) Expression of CD80 (left) and CD86 (right) in HLA-A2+ CD11c^+^ DCs, relative to DCs cultured with untransduced Tregs (dotted line). (**D**) DCs suppression assays performed with or without 10 μg/mL CTLA4-Ig. For C&D, data are shown relative to DCs cultured with untransduced Tregs which were normalized to 100% (dotted lines). Data shown as mean±SEM and pooled from 5 (**C**) and 2 (**D**) independent experiments. Statistical significance was determined using one-way ANOVA with a Holm-Sidak post-test, *p<0.05, **p<0.01, ***p<0.001, ****p<0.0001.

## DISCUSSION

Understanding how the structure of a CAR affects Treg function is critical to guide their clinical implementation. Here we studied how different CAR co-stimulatory domains affect Treg function in an immunocompetent mouse model of skin transplantation. Whilst 4-1BB- and OX40-encoding CAR-Tregs did not have a significant therapeutic effect, CD28-, ICOS-, PD1- and GITR-encoding CAR-Tregs were similarly efficacious in vivo. Further comparisons between Tregs expressing a first (A2.ζ) or second (A2.28ζ) generation CAR revealed equivalent function, leading us to study a possible role for co-stimulation via the native CD28 receptor. These studies showed that native CD28 signaling can compensate for a lack of CAR-mediated co-stimulation, providing a significant advance in our understanding of how CAR-Tregs interact with host immune cells and regulate alloimmunity.

We and others previously compared the function of CARs encoding different co-stimulatory domains in Tregs using immunodeficient mouse models and found that a CD28 co-stimulatory domain was optimal for Treg potency, stability, persistence and in vivo function (10, 11, 16, 48). CARs carrying alternative co-stimulatory domains had limited in vitro and in vivo function (10, 11, 16, 48). In contrast, in the immunocompetent transplant setting used here, Tregs expressing CARs encoding co-stimulatory domains from ICOS, PD1 or GITR were similar to CD28 in terms of protection from skin rejection (although not DSA generation). This finding was particularly intriguing for the PD1-encoding CAR, since it only weakly activates T cells (46) and Tregs (10). Our data suggest that, at least for some CARs, this could be related to the combination of native CD28 and CAR-mediated signaling, with the former compensating for lower CAR-mediated activation.

CD28 is a major co-stimulatory receptor for Tregs (50-53), but these cells express a variety of costimulatory molecules (54, 55) that have positive or negative effects (54, 56). Thus, it is possible that certain combinations of co-stimulatory signaling driven by the CAR and/or natural co-receptors could be harmful and cause Treg dysfunction(48). Indeed, in previous studies using immunodeficient mouse models, CARs carrying co-stimulatory domains from TNFR family members, such as 41BB, showed no therapeutic protection (10, 11, 16, 48) and were associated with exhaustion (48) and loss of Treg stability (10, 48). Similarly, in our immunocompetent transplant setting, CARs encoding costimulatory domains from 41BB and OX40 showed no protection from graft rejection or control of DSAs generation. However, there were also no signs of tonic signaling or loss of Foxp3 or Helios expression even after 4 weeks after adoptive transfer. Thus, in vivo co-stimulation from the CD28 receptor might overcome the deleterious effect of these otherwise harmful co-stimulatory signals (57).

CARs were originally developed for use in cancer with the goal of directing T cells to kill tumor cells (58). These tumor cells often overexpress coinhibitory receptors as an immune escape mechanism and may not express costimulatory molecules such as CD80 or CD86 (59, 60). As such, first-generation CARs lacking co-stimulation showed modest outcomes in a cancer setting (61-63) and the provision of co-stimulation in a second-generation CAR format greatly increased their clinical success (64). In Tregs, studies of first-generation CARs are limited, with only one study in immunodeficient mice showing little protection from xenogeneic GvHD (10). Conversely, we found that in an immunocompetent mouse setting, first- and second-generation CAR-Tregs offer the same protection from rejection. Distinct from CAR T-cells, APCs are a major target for Treg suppression (44), leading us to speculate that first-generation CAR-Tregs could receive natural co-stimulation via these cells. This possibility is supported by our in vitro data with Tregs showing that native CD28 co-stimulation compensates for absent CAR-encoded co-stimulation. Similar findings were reported with CAR-T cells in vitro (65, 66): if co-stimulatory molecules are provided, first- and second-generation CARs equivalently activate T cells. Collectively these data highlight that in vivo, CAR-Treg function is ultimately determined by an integrated response to CAR- and native co-stimulation-mediated signaling.

An outstanding question is where would CAR-Tregs encounter donor antigen and co-stimulation? Skin-resident APCs play an important role in the regulation of alloimmunity (67-70) and in our immunocompetent skin graft model, these cells could deliver both CAR and co-stimulatory signals to CAR-Tregs. Skin donor APCs could also migrate to surrounding lymphoid nodes (LN) (71) and/or host APCs could be cross-dressed with HLA-A2 via exosome-mediated mechanisms (72-74). The fact that first and CD28-based second-generation CAR-Tregs were both able to control DSA generation suggests that CAR-Tregs migrating to LNs not only receive CAR signals but also co-stimulation.

Notably, CAR-Treg therapy delayed graft rejection but did not induce indefinite graft survival. Similar results have been reported by previous studies using different models of transplantation (8, 9). A potential reason for this could be the inability of CAR-Tregs to control the high numbers of alloreactive T cells generated after transplantation, an issue that can be resolved by administrating cytotoxic or immunosuppressive preconditioning treatments before infusing Tregs (75, 76). Another reason could be the high stringency of immunocompetent transplant mouse models of transplantation, particularly of the skin allograft model. Studies exploring the use of A2-CAR Tregs alone in a single HLA-A2-mismatched heart transplant model also failed to induce long-term tolerance but extended graft protection longer than our skin allograft model (17). However, skin allograft models facilitate testing of multiple CAR-Treg groups in parallel which is not feasible with other less-stringent transplant models due to their complexity.

Another potential factor limiting for CAR-Tregs is their short in vivo persistence, which could be driven by multiple mechanisms. CAR-Tregs uptake HLA-A2 molecules by trogocytosis (15), thus the resulting HLA-A2^+^ CAR-Tregs could become targets for anti-HLA-A2 Abs and be depleted. Low IL-2 levels of IL-2 could impact CAR-Treg persistence (77), as could diminishing levels of the target antigen which may become limiting as rejection progresses. A consideration is that the persistence data reported here were from blood so may not capture cells which have re-localized to tissues. Overall, investigation of strategies to enhance persistence, such as by repeat dosing, or co-administration of adjunct therapies such as IL-2, is warranted.

Overall, our results contribute to the understanding of how alternative co-stimulatory domains impact the in vivo function of CAR-Tregs and demonstrate that CAR-mediated co-stimulation in Tregs is not essential for in vivo function. These data provide an important step forward in our understanding of the biology of CAR-Tregs and how to best optimize them for clinical applications.

## METHODS

### Generation of signaling domain CAR variants

CAR variants were generated by replacing the transmembrane and co-stimulatory domains of a previously characterized A2-specific CAR (10, 15). Domain sequences were obtained from Uniprot and codon optimized for mouse (**Supplemental Table 2**). The resulting CARs encoded an A2-specific scFv (12), a CD8α-derived hinge, a c-Myc epitope tag, the indicated transmembrane and co-stimulatory domains, and CD3ζ. A HER2-specific CAR served as an antigen-non-specific negative control (10, 15). CARs were cloned into an MSCV-based vector upstream of an IRES-monomeric Kusabira-Orange2 (mKO2) reporter and retroviral particles were generated as described (15).

### Animals

Bl/6, Bl/6-Foxp3^gfp^ × Thy1.1 Bl/6, HLA-A2^+^ Bl/6 mice (B6.Cg-Tg(HLA-A/H2-D)2Enge/J) and OTII Bl6 mice (B6.Cg-Tg(TcraTcrb)425Cbn/J) were purchased from Jackson Laboratories and bred in-house under specific pathogen-free conditions.

### CAR-Treg generation

CAR-Tregs were generated as described (15, 78). Briefly, lymph nodes and spleen from 16-24-week-old female or male C57Bl/6-Foxp3^gfp^ × Thy1.1 mice were collected and CD4^+^ T cells isolated by negative selection (STEMCELL Technologies). Tregs were sorted as live CD4^+^CD8^-^Thy1.1^+^Foxp3^gfp+^ using a MoFlow® Astrios (Beckman Coulter) (**Supplemental Figure 1A**), stimulated with anti-CD3/CD28 dynabeads (ThermoFisher Scientific), expanded in the presence of recombinant human IL-2 (1000 U/mL, Proleukin) and rapamycin (50 nmol/L, Sigma-Aldrich), and transduced after 2 days. Dynabeads were removed on day 7 and cells were rested overnight in 1000U/mL, or 100U/mL for 2 days, prior to use for in vivo or in vitro assays, respectively. CAR expression and Treg purity were determined after expansion (**Supplemental Figure 1B**).

### Proliferation, activation and cytokine production

CAR-Tregs were labelled with CPDeFluor450 proliferation dye (eBioscience), then stimulated with irradiated (125 Gy) HLA-A2^pos^CD86^neg^, HLA-A2^pos^CD86^pos^ or HLA-A2^neg^CD86^neg^ K562 cells at a 1:2 (K562:Treg) ratio with 100 U/mL IL-2. After 72 hrs activation markers and proliferation (CPDe450 dilution or Ki67 expression) were assessed by flow cytometry and cell culture supernatants were collected to measure cytokine secretion using a cytometric bead array (BD Biosciences). Where stated, CTLA4-Ig (Orencia) and/or an anti-CD28 agonist antibody (Clone:37.51, BD bioscience) were added at 10 μg/mL.

### Suppression assays

For T cell suppression, responder CD4^+^ T cells were isolated from OTII BL/6 mice by negative selection (STEMCELL Technologies). Splenocytes from wild-type or HLA-A2^+^ BL/6 mice were depleted of Thy1.2^+^ cells (STEMCELL Technologies), irradiated (20 Gy) and 175,000 were co-cultured with 25,000 OTII T cells with 200 ng/mL OVA_323-339_ peptide (Sigma-Aldrich) and varying ratios of Tregs. OTII proliferation was measured by flow cytometry after 4 days and % suppression was calculated as the inhibition of Tresponder proliferation, relative to Tresponders cultured without Tregs.

For DC suppression, splenic CD11c^+^ DCs were isolated by positive selection (STEMCELL Technologies) from wild-type or HLA-A2^+^ BL/6 mice and cultured with CAR-Tregs (1:2 or 1:5 DC:Treg ratio). Suppressive effects of CAR-Tregs were measured as % decreased expression of costimulatory (CD80 and CD86) and MHC-II molecules on DCs after 1 and/or 2 days.

### Skin transplantation

10-14-week-old female and male wild-type C57BL/6 mice were transplanted with dorsal skin grafts from sex-matched, wild-type or HLA-A2^+^ BL/6 mice. Where stated, mice were injected with 1×10^6^ CAR-Tregs (equivalent to 30-50×10^6^/kg) into the tail vein at the time of transplantation (15). Grafts were covered with a petroleum jelly gauze patch and wrapped with CoFlex bandage (3M, Nexcare). Bandages were removed after 10-days and grafts monitored for rejection until 30 days post-transplantation. Graft rejection was defined as described(15). Peripheral blood and plasma were collected weekly to track CAR-Tregs and antibodies. Red blood cells were lysed using ammonium chloride and Fc receptors were blocked using anti-mouse CD16/CD32 (BD Bioscience) before staining.

### Anti-HLA-A2 IgG quantification

Anti-HLA-A2 IgG titers were determined using a cell-based ELISA. HLA-A2^pos^ K562 and control K562 cells were seeded in a 96-well plate and blocked with rat serum (STEMCELL) for 30 mins at RT. Plasma samples were added (1:800 dilution) and incubated for 1 hour at RT. A goat anti-mouse IgG APC secondary antibody (Invitrogen) was added (1:700 dilution) and incubated for 1 hour at RT. A standard curve was made using purified anti-HLA-A2 antibody (BD, clone:BB7.2). Cells were analyzed by flow cytometry and concentration was calculated based on MFI using a 4PL curve.

### Flow cytometry

Flow cytometry was performed in adherence to “Guidelines for the use of flow cytometry and cell sorting in immunological studies (third edition)” (79). Flow cytometric antibodies are shown in **Supplemental Table 3**. Cells were extracellularly stained in presence of Fixable Viability Dye (FVD) eFluor™ 780 (ThermoFisher Scientific) to exclude dead cells.

Staining for intracellular markers was performed using the Foxp3/Transcription Factor Staining Buffer Set (Thermo Fisher Scientific). Data were acquired using an LSR Fortessa II, A5 Symphony (BD Biosciences) or CytoFLEX (Beckman Coulter), and analyzed using FlowJo version 10.7.1 (Tree Star).

### Statistics

Data were analyzed using GraphPad Prism 9.3.1 and are shown as mean±SEM. Statistical significance were determined using Pearson correlation, one-way and two-way analysis of variance (ANOVA) with a Holm-Sidak post-test or by log-rank (Mantel-Cox) test for survival curve comparisons.

### Study Approval

Animal experiments were approved by the University of British Columbia Animal Care and Use Committee (A19-0136).

## Acknowledgements

Supported by grants from the US Department of Defense (W81XWH-19-1-0351) and the Canadian Institutes for Health Research (FDN-154304). IRS and DAB are supported by salary awards from the Canadian Institutes for Health Research and Michael Smith Health Research BC. MKL receives a salary award from the BC Children’s Hospital Research Institute and is a Canada Research Chair in Engineered Immune Tolerance. Some figures were created with BioRender (http://biorender.com)

## REFERENCES

1. Lam AJ, Hoeppli RE, and Levings MK. Harnessing Advances in T Regulatory Cell Biology for Cellular Therapy in Transplantation. Transplantation. 2017;101(10):2277–87.

2. Raffin C, Vo LT, and Bluestone JA. Treg cell-based therapies: challenges and perspectives. Nature Reviews Immunology. 2020;20(3):158–72.

3. Ferreira LMR, Muller YD, Bluestone JA, and Tang Q. Next-generation regulatory T cell therapy. Nature Reviews Drug Discovery. 2019;18(10):749–69.

4. Sawitzki B, Harden PN, Reinke P, Moreau A, Hutchinson JA, Game DS, et al. Regulatory cell therapy in kidney transplantation (The ONE Study): a harmonised design and analysis of seven non-randomised, single-arm, phase 1/2A trials. Lancet. 2020;395(10237):1627–39.

5. Harden PN, Game DS, Sawitzki B, Van der Net JB, Hester J, Bushell A, et al. Feasibility, long-term safety, and immune monitoring of regulatory T cell therapy in living donor kidney transplant recipients. Am J Transplant. 2021;21(4):1603–11.

6. Sánchez-Fueyo A, Whitehouse G, Grageda N, Cramp ME, Lim TY, Romano M, et al. Applicability, safety, and biological activity of regulatory T cell therapy in liver transplantation. Am J Transplant. 2020;20(4):1125–36.

7. Roemhild A, Otto NM, Moll G, Abou-El-Enein M, Kaiser D, Bold G, et al. Regulatory T cells for minimising immune suppression in kidney transplantation: phase I/IIa clinical trial. BMJ. 2020;371:m3734.

8. Rosado-Sánchez I, and Levings MK. Building a CAR-Treg: Going from the basic to the luxury model. Cellular Immunology. 2020;358(September).

9. Dawson NAJ, Vent-Schmidt J, and Levings MK. Engineered tolerance: Tailoring development, function, and antigen-specificity of regulatory T cells. Frontiers in Immunology. 2017;8.

10. Dawson NAJ, Rosado-Sánchez I, Novakovsky GE, Fung VCW, Huang Q, McIver E, et al. Functional effects of chimeric antigen receptor co-receptor signaling domains in human Tregs. Sci Transl Med. 2020;12(557).

11. Imura Y, Ando M, Kondo T, Ito M, and Yoshimura A. CD19-targeted CAR regulatory T cells suppress B cell pathology without GvHD. JCI Insight. 2020;5(14).

12. MacDonald KG, Hoeppli RE, Huang Q, Gillies J, Luciani DS, Orban PC, et al. Alloantigen-specific regulatory T cells generated with a chimeric antigen receptor. Journal of Clinical Investigation. 2016;126(4):1413–24.

13. Boardman DA, Philippeos C, Fruhwirth GO, Ibrahim MAA, Hannen RF, Cooper D, et al. Expression of a Chimeric Antigen Receptor Specific for Donor HLA Class I Enhances the Potency of Human Regulatory T Cells in Preventing Human Skin Transplant Rejection. American Journal of Transplantation. 2017;17(4):931–43.

14. Noyan F, Zimmermann K, Hardtke-Wolenski M, Knoefel A, Schulde E, Geffers R, et al. Prevention of Allograft Rejection by Use of Regulatory T Cells With an MHC-Specific Chimeric Antigen Receptor. American Journal of Transplantation. 2017;17(4):917–30.

15. Sicard A, Lamarche C, Speck M, Wong M, Rosado-Sánchez I, Blois M, et al. Donor-specific chimeric antigen receptor Tregs limit rejection in naive but not sensitized allograft recipients. American Journal of Transplantation. 2020(January):1–12.

16. Boroughs AC, Larson RC, Choi BD, Bouffard AA, Riley LS, Schiferle E, et al. Chimeric antigen receptor costimulation domains modulate human regulatory T cell function. JCI Insight. 2019;5(8).

17. Wagner JC, Ronin E, Ho P, Peng Y, and Tang Q. Anti-HLA-A2-CAR Tregs prolong vascularized mouse heterotopic heart allograft survival. Am J Transplant. 2022;22(9):2237–45.

18. Schreeb K, Culme-Seymour E, Ridha E, Dumont C, Atkinson G, Hsu B, et al. Study Design: Human Leukocyte Antigen Class I Molecule A. Kidney Int Rep. 2022;7(6):1258–67.

19. Guedan S, Calderon H, Posey AD, and Maus MV. Engineering and Design of Chimeric Antigen Receptors. Molecular Therapy - Methods and Clinical Development. 2018;12:145–56.

20. Rafiq S, Hackett CS, and Brentjens RJ. Engineering strategies to overcome the current roadblocks in CAR T cell therapy. Nature Reviews Clinical Oncology. 2020;17(3):147–67.

21. Boardman DA, and Levings MK. Emerging strategies for treating autoimmune disorders with genetically modified Treg cells. J Allergy Clin Immunol. 2022;149(1):1–11.

22. Søndergaard H, Kvist PH, and Haase C. Human T cells depend on functional calcineurin, tumour necrosis factor-α and CD80/CD86 for expansion and activation in mice. Clin Exp Immunol. 2013;172(2):300–10.

23. Walsh NC, Kenney LL, Jangalwe S, Aryee KE, Greiner DL, Brehm MA, et al. Humanized Mouse Models of Clinical Disease. Annu Rev Pathol. 2017;12:187–215.

24. Kenney LL, Shultz LD, Greiner DL, and Brehm MA. Humanized Mouse Models for Transplant Immunology. Am J Transplant. 2016;16(2):389–97.

25. Shultz LD, Keck J, Burzenski L, Jangalwe S, Vaidya S, Greiner DL, et al. Humanized mouse models of immunological diseases and precision medicine. Mamm Genome. 2019;30(5-6):123–42.

26. Vignali DAA, Collison LW, and Workman CJ. How regulatory T cells work. Nature Reviews Immunology. 2008;8(7):523–32.

27. Qureshi OSZ, Yong Nakamura K, Attridge K, Manzotti C, Schmidt EM, Baker J, et al. Trans-endocytosis of CD80 and CD86: A molecular basis for the cell-extrinsic function of CTLA-4. Science. 2011;332(6029):600–3.

28. Tekguc M, Wing JB, Osaki M, Long J, and Sakaguchi S. Treg-expressed CTLA-4 depletes CD80/CD86 by trogocytosis, releasing free PD-L1 on antigen-presenting cells. Proc Natl Acad Sci U S A. 2021;118(30).

29. Liang B, Workman C, Lee J, Chew C, Dale BM, Colonna L, et al. Regulatory T cells inhibit dendritic cells by lymphocyte activation gene-3 engagement of MHC class II. J Immunol. 2008;180(9):5916–26.

30. Akkaya B, Oya Y, Akkaya M, Al Souz J, Holstein AH, Kamenyeva O, et al. Regulatory T cells mediate specific suppression by depleting peptide-MHC class II from dendritic cells. Nat Immunol. 2019;20(2):218–31.

31. Veldhoen M, Moncrieffe H, Hocking RJ, Atkins CJ, and Stockinger B. Modulation of dendritic cell function by naive and regulatory CD4+ T cells. J Immunol. 2006;176(10):6202–10.

32. Iikuni N, Lourenço EV, Hahn BH, and La Cava A. Cutting edge: Regulatory T cells directly suppress B cells in systemic lupus erythematosus. J Immunol. 2009;183(3):1518–22.

33. Weingartner E, and Golding A. Direct control of B cells by Tregs: An opportunity for long-term modulation of the humoral response. Cell Immunol. 2017;318:8–16.

34. Wollenberg I, Agua-Doce A, Hernández A, Almeida C, Oliveira VG, Faro J, et al. Regulation of the germinal center reaction by Foxp3+ follicular regulatory T cells. J Immunol. 2011;187(9):4553–60.

35. Linterman MA, Pierson W, Lee SK, Kallies A, Kawamoto S, Rayner TF, et al. Foxp3+ follicular regulatory T cells control the germinal center response. Nat Med. 2011;17(8):975–82.

36. Wing JB, Ise W, Kurosaki T, and Sakaguchi S. Regulatory T cells control antigen-specific expansion of Tfh cell number and humoral immune responses via the coreceptor CTLA-4. Immunity. 2014;41(6):1013–25.

37. Sage PT, Paterson AM, Lovitch SB, and Sharpe AH. The coinhibitory receptor CTLA-4 controls B cell responses by modulating T follicular helper, T follicular regulatory, and T regulatory cells. Immunity. 2014;41(6):1026–39.

38. Wang CJ, Heuts F, Ovcinnikovs V, Wardzinski L, Bowers C, Schmidt EM, et al. CTLA-4 controls follicular helper T-cell differentiation by regulating the strength of CD28 engagement. Proc Natl Acad Sci U S A. 2015;112(2):524–9.

39. Alegre ML, Lakkis FG, and Morelli AE. Antigen Presentation in Transplantation. Trends Immunol. 2016;37(12):831–43.

40. Chakraverty R, and Sykes M. The role of antigen-presenting cells in triggering graft-versus-host disease and graft-versus-leukemia. Blood. 2007;110(1):9–17.

41. Lei YM, Sepulveda M, Chen L, Wang Y, Pirozzolo I, Theriault B, et al. Skin-restricted commensal colonization accelerates skin graft rejection. JCI Insight. 2019;5.

42. Loupy A, and Lefaucheur C. Antibody-Mediated Rejection of Solid-Organ Allografts. N Engl J Med. 2018;379(12):1150–60.

43. Loupy A, Hill GS, and Jordan SC. The impact of donor-specific anti-HLA antibodies on late kidney allograft failure. Nat Rev Nephrol. 2012;8(6):348–57.

44. Wardell M Christine MNK, Levings K Megan, Cook Laura. Cross talk between human regulatory T cells and antigen-presenting cells: Lessons for clinical applications. Eur J Immunol. 2021;51:27–38.

45. Wing JB, Tay C, and Sakaguchi S. Control of Regulatory T Cells by Co-signal Molecules. Adv Exp Med Biol. 2019;1189:179–210.

46. Fedorov VD, Themeli M, and Sadelain M. PD-1– and CTLA-4–Based Inhibitory Chimeric Antigen Receptors (iCARs) Divert Off-Target Immunotherapy Responses. Sci Transl Med. 2013;11(5).

47. Guedan S, Posey AD, Shaw C, Wing A, Da T, Patel PR, et al. Enhancing CAR T cell persistence through ICOS and 4-1BB costimulation. JCI insight. 2018;3(1).

48. Lamarthee B, Marchal A, Charbonnier S, Blein T, Leon J, Martin E, et al. Transient mTOR inhibition rescues 4-1BB CAR-Tregs from tonic signal-induced dysfunction. Nat Commun. 2021;12(1):6446.

49. Muller YD, Nguyen DP, Ferreira LMR, Ho P, Raffin C, Valencia RVB, et al. The CD28-Transmembrane Domain Mediates Chimeric Antigen Receptor Heterodimerization With CD28. Front Immunol. 2021;12:639818.

50. Hombach AA, Kofler D, Hombach A, Rappl G, and Abken H. Effective proliferation of human regulatory T cells requires a strong costimulatory CD28 signal that cannot be substituted by IL-2. J Immunol. 2007;179(11):7924–31.

51. Tang Q, Henriksen KJ, Boden EK, Tooley AJ, Ye J, Subudhi SK, et al. Cutting edge: CD28 controls peripheral homeostasis of CD4+CD25+ regulatory T cells. J Immunol. 2003;171(7):3348–52.

52. Gogishvili T, Lühder F, Goebbels S, Beer-Hammer S, Pfeffer K, and Hünig T. Cell-intrinsic and -extrinsic control of Treg-cell homeostasis and function revealed by induced CD28 deletion. Eur J Immunol. 2013;43(1):188–93.

53. Golovina TN, Mikheeva T, Suhoski MM, Aqui NA, Tai VC, Shan X, et al. CD28 costimulation is essential for human T regulatory expansion and function. J Immunol. 2008;181(4):2855–68.

54. Wing JB, Tay C, and Sakaguchi S. Control of Regulatory T Cells by Co-signal Molecules. Adv Exp Med Biol. 2019;1189:179–210.

55. Hoeppli RE, Wu D, Cook L, and Levings MK. The environment of regulatory T cell biology: cytokines, metabolites, and the microbiome. Front Immunol. 2015;6:61.

56. Bour-Jordan H, and Bluestone JA. Regulating the regulators: Costimulatory signals control the homeostasis and function of regulatory T cells. Immunological Reviews. 2009;229(1):41–66.

57. Choi BK, Bae JS, Choi EM, Kang WJ, Sakaguchi S, Vinay DS, et al. 4-1BB-dependent inhibition of immunosuppression by activated CD4+CD25+ T cells. J Leukoc Biol. 2004;75(5):785–91.

58. Hou AJ, Chen LC, and Chen YY. Navigating CAR-T cells through the solid-tumour microenvironment. Nat Rev Drug Discov. 2021;20(7):531–50.

59. Capece D, Verzella D, Fischietti M, Zazzeroni F, and Alesse E. Targeting costimulatory molecules to improve antitumor immunity. J Biomed Biotechnol. 2012;2012:926321.

60. Zou W, and Chen L. Inhibitory B7-family molecules in the tumour microenvironment. Nat Rev Immunol. 2008;8(6):467–77.

61. Eshhar Z, Waks T, Gross G, and Schindler DG. Specific activation and targeting of cytotoxic lymphocytes through chimeric single chains consisting of antibody-binding domains and the gamma or zeta subunits of the immunoglobulin and T-cell receptors. Proc Natl Acad Sci U S A. 1993;90(2):720–4.

62. Brocker T, and Karjalainen K. Signals through T cell receptor-zeta chain alone are insufficient to prime resting T lymphocytes. J Exp Med. 1995;181(5):1653–9.

63. Brocker T. Chimeric Fv-zeta or Fv-epsilon receptors are not sufficient to induce activation or cytokine production in peripheral T cells. Blood. 2000.

64. Sterner RC, and Sterner RM. CAR-T cell therapy: current limitations and potential strategies. Blood Cancer J. 2021;11(4):69.

65. Zhong XS, Matsushita M, Plotkin J, Riviere I, and Sadelain M. Chimeric antigen receptors combining 4-1BB and CD28 signaling domains augment PI 3 kinase/AKT/Bcl-X L activation and CD8 T cell-mediated tumor eradication. Molecular Therapy. 2010;18(2):413–20.

66. Friedmann-Morvinski D, Bendavid A, Waks T, Schindler D, and Eshhar Z. Redirected primary T cells harboring a chimeric receptor require costimulation for their antigen-specific activation. Blood. 2005;105(8):3087–93.

67. Kashem SW, Haniffa M, and Kaplan DH. Antigen-Presenting Cells in the Skin. Annu Rev Immunol. 2017;35:469–99.

68. Scheib N, Tiemann J, Becker C, Probst HC, Raker VK, and Steinbrink K. The Dendritic Cell Dilemma in the Skin: Between Tolerance and Immunity. Front Immunol. 2022;13:929000.

69. Benichou G, Yamada Y, Yun SH, Lin C, Fray M, and Tocco G. Immune recognition and rejection of allogeneic skin grafts. Immunotherapy. 2011;3(6):757–70.

70. Debes GF, and McGettigan SE. Skin-Associated B Cells in Health and Inflammation. J Immunol. 2019;202(6):1659–66.

71. Morelli AE. Dendritic cells of myeloid lineage: the masterminds behind acute allograft rejection. Curr Opin Organ Transplant. 2014;19(1):20–7.

72. Liu Q, Rojas-Canales DM, Divito SJ, Shufesky WJ, Stolz DB, Erdos G, et al. Donor dendritic cell-derived exosomes promote allograft-targeting immune response. J Clin Invest. 2016;126(8):2805–20.

73. Prunevieille A, Babiker-Mohamed MH, Aslami C, Gonzalez-Nolasco B, Mooney N, and Benichou G. T cell antigenicity and immunogenicity of allogeneic exosomes. Am J Transplant. 2021;21(7):2583–9.

74. Zeng F, Chen Z, Chen R, Shufesky WJ, Bandyopadhyay M, Camirand G, et al. Graft-derived extracellular vesicles transported across subcapsular sinus macrophages elicit B cell alloimmunity after transplantation. Sci Transl Med. 2021;13(585).

75. Lee K, Nguyen V, Lee KM, Kang SM, and Tang Q. Attenuation of donor-reactive T cells allows effective control of allograft rejection using regulatory T cell therapy. Am J Transplant. 2014;14(1):27–38.

76. Tsang JY, Tanriver Y, Jiang S, Xue SA, Ratnasothy K, Chen D, et al. Conferring indirect allospecificity on CD4+CD25+ Tregs by TCR gene transfer favors transplantation tolerance in mice. J Clin Invest. 2008;118(11):3619–28.

77. Ratnasothy K, Jacob J, Tung S, Boardman D, Lechler RI, Sanchez-Fueyo A, et al. IL-2 therapy preferentially expands adoptively transferred donor-specific Tregs improving skin allograft survival. Am J Transplant. 2019;19(7):2092–100.

78. Wu D, Wong MQ, Vent-Schmidt J, Boardman DA, Steiner TS, and Levings MK. A method for expansion and retroviral transduction of mouse regulatory T cells. J Immunol Methods. 2021;488:112931.

79. Cossarizza A, Chang HD, Radbruch A, Abrignani S, Addo R, Akdis M, et al. Guidelines for the use of flow cytometry and cell sorting in immunological studies (third edition). Eur J Immunol. 2021;51(12):2708–3145.

