## Supplemental Data for "Integration of exogenous and endogenous co-stimulatory signals by CAR-Tregs"

### SUPPLEMENTARY DATA

**Supplemental Figure 1**

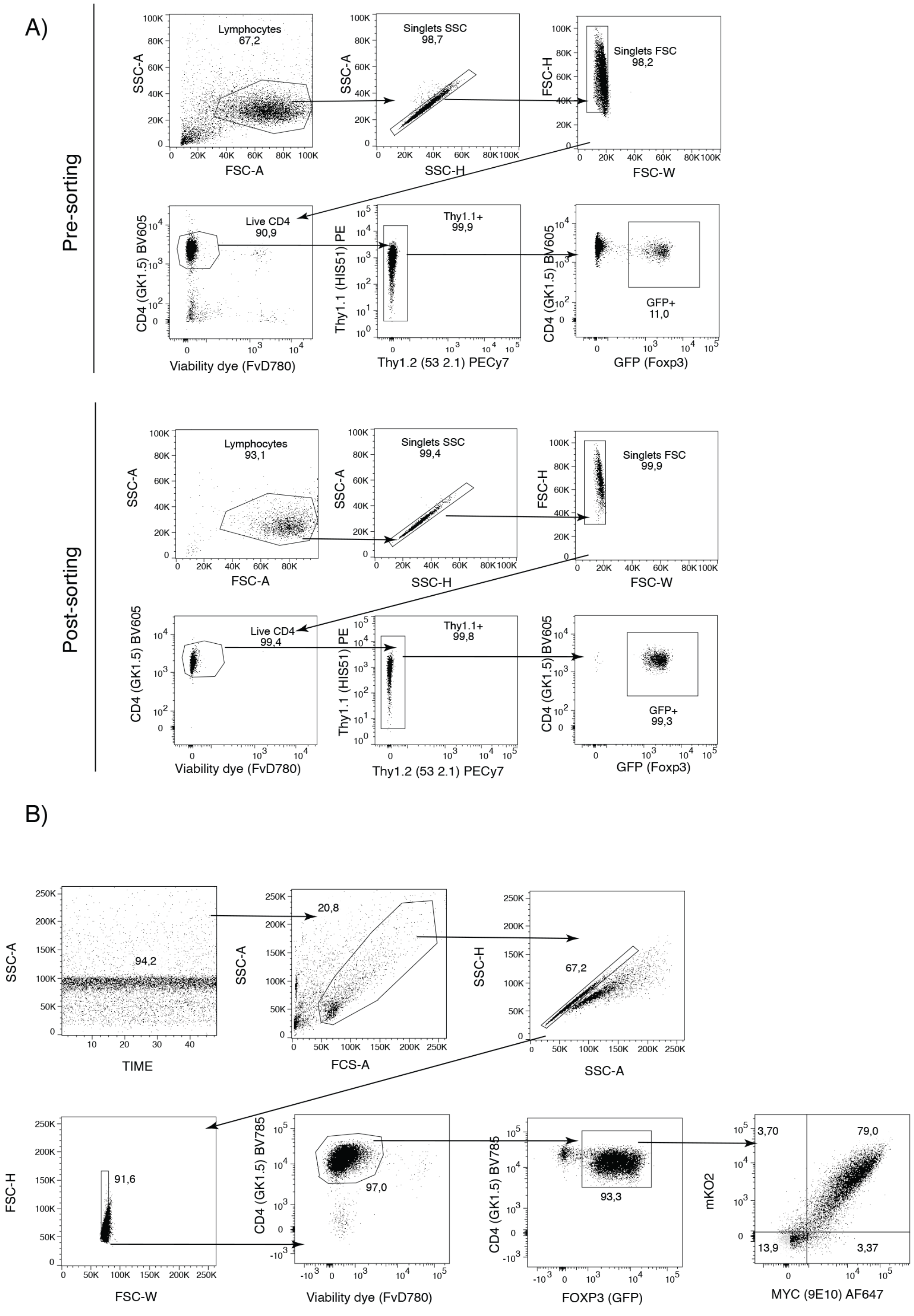


**Supplementary Figure 1**. ***Sorting and purity check flow panels.*** (**A**) Representative example of the pre- and post-sorting flow panel used to isolate Tregs (defined as live CD4+CD8-

Thy1.1+Foxp3$^{gfp}$+ cells). (**B**) Representative example of the flow panel used to check Treg purity and CAR-expression after Treg expansion at day 7.

**Supplemental Figure 2**

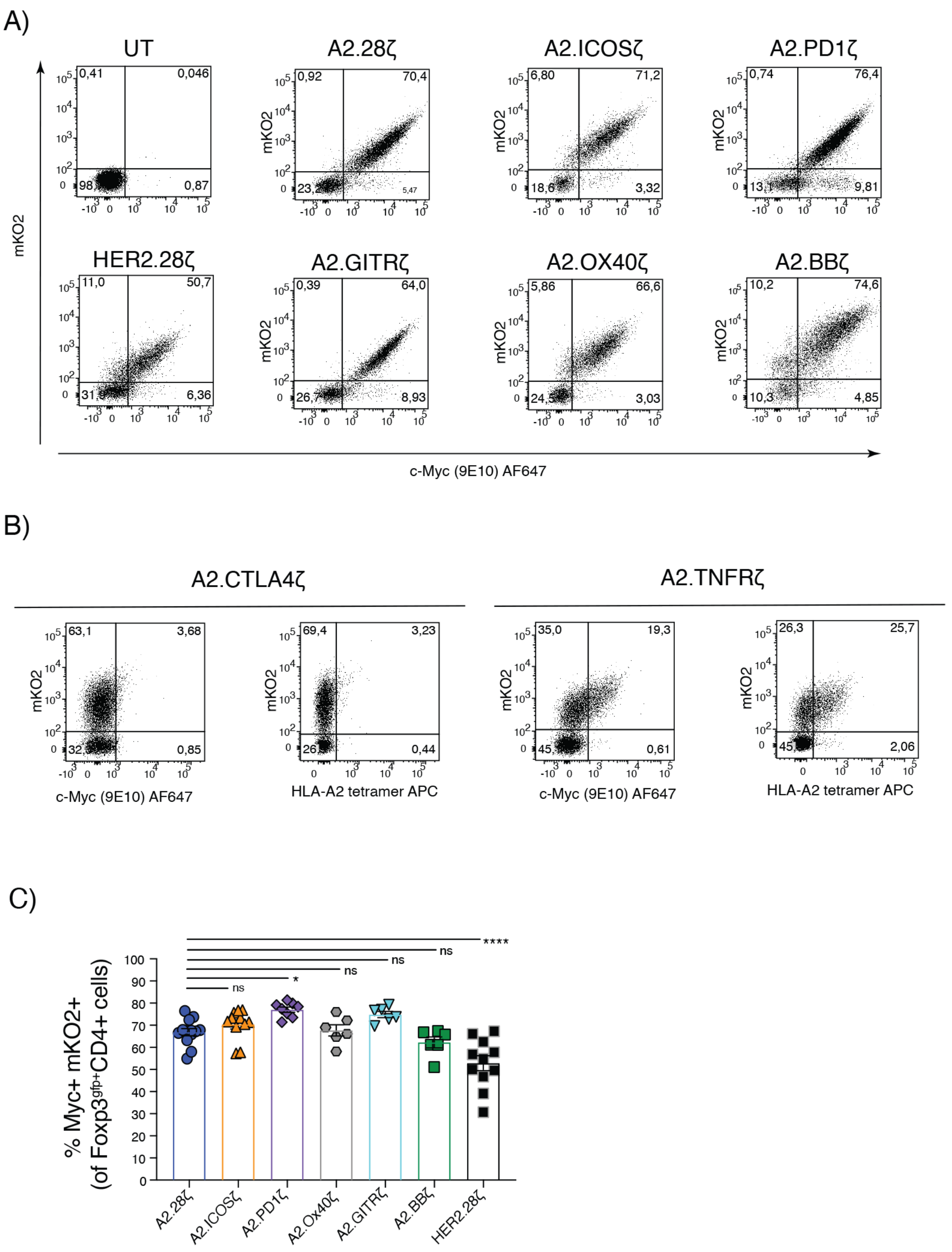


**Supplementary Figure 2**: *Co-stimulatory domain variant CAR expression:* (**A**) Representative examples of the expression of co-stimulatory domain CAR variants defined by c-Myc and mKO2 co-expression in Tregs. (**B**) Lack of CAR expression or binding to HLA-A2 for the CTLA4- and TNFR2-based CAR variants. (**C**) Frequencies of c-Myc and mKO2 co-expression in Tregs, gated in live CD4+foxp3$^{gfp}$+ cells. Data pooled from at least 6 independent experiments and shown as mean±SEM. Statistical significance was determined using one-way ANOVA with a Holm-Sidak post-test, *p<0.05, **p<0.01, ***p<0.001, ****p<0.0001.

**Supplemental Figure 3**

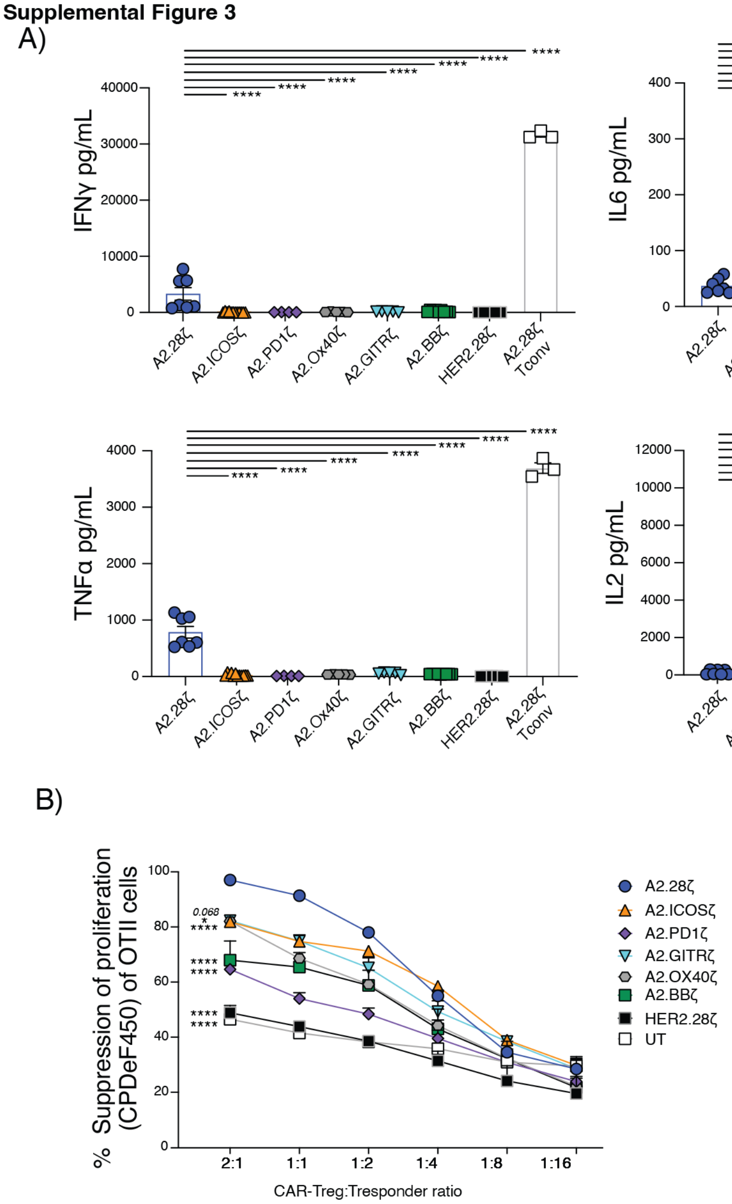


**Supplementary Figure 3:** *Supplementary data for the in vitro characterization of co-stimulatory domain CAR variants.* (**A**) Cytokine production of CAR-Tregs cultured for 3-days with HLA-A2$^+$ K562 cells. Data pooled from 3 independent experiments. (**B**) CAR-Treg suppression of the OTII CD4 T responder proliferation, as determined by CPDeF450 dilution. Data pooled from 2 independent experiments. UT = Untransduced. Data pooled from 4 independent experiments. Data shown as mean±SEM. Statistical significance was determined using one-way (**A**) two-way (**B**) ANOVA with a Holm-Sidak post-test. *p<0.05, **p<0.01, ***p<0.001. ****p<0.0001.

**Supplemental Figure 4**

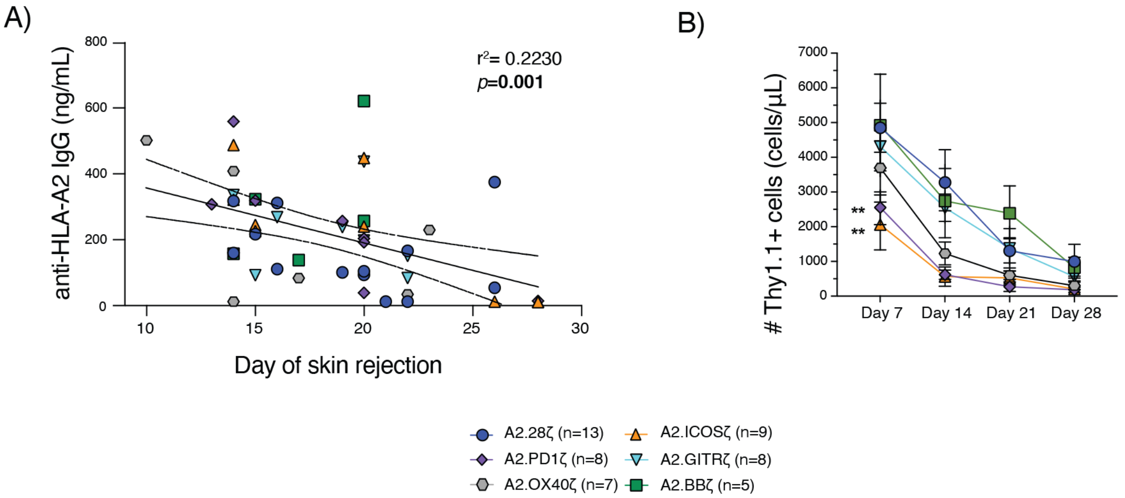


**Supplementary Figure 4:** *Supplementary data for the in vivo characterization of co-stimulatory domain CAR variants.* (**A**) Correlation between levels of anti-HLA-A2 IgG Abs in plasma at day 14 and skin graft rejection date of mice receiving Tregs bearing CD28 and TNFR family-based co-stimulatory domain CARs. (**B**) Count of CAR-Tregs in peripheral blood of mice transplanted with HLA-A2-expressing skin and intravenously administered with $1x10^6$ CAR-Tregs. Data pooled from 4 independent experiments. Data shown as mean±SEM (**B**). Statistical significance was determined using Pearson Correlation (**A**) or two-way ANOVA (**B**) with a Holm-Sidak post-test. *p<0.05, **p<0.01, ***p<0.001. ****p<0.0001.

**Supplemental Figure 5**

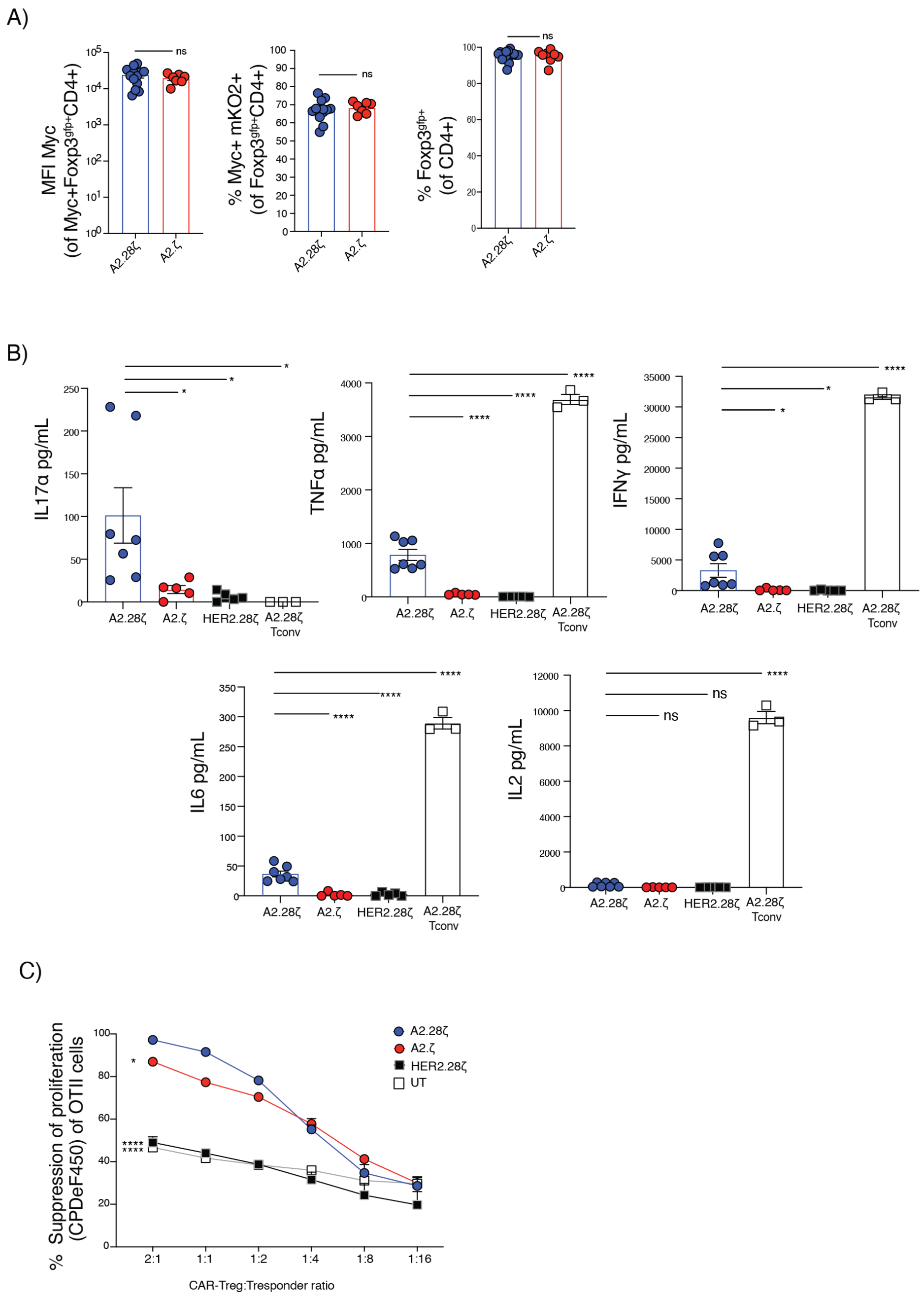


**Supplementary Figure 5:** *Supplementary material for the in vitro comparison of first- and second-generation CARs*: **A**) MFI of CAR expression, co-expression of Myc and mKO2 and Foxp3$^{gfp}$ expression for first- and second-generation CAR-Tregs after expansion, gated on total live Myc+CD4$^{+}$Foxp3$^{gfp+}$ or live CD4$^{+}$ cells. Data pooled from at least 6 independent experiments. (**B**) Cytokine production of CAR-Tregs cultured for 3-days with HLA-A2$^{pos}$ K562 cells. Data pooled from 3 independent experiments (**C**) CAR-Treg suppression of the OTII CD4 T responder proliferation, as determined by CPDeF450 dilution. Data pooled from 2 independent experiments UT = Untransduced. Data shown as mean±SEM. Statistical significance was determined using (**A**) t-student, one-way (**B,C**) two-way ANOVA with a Holm-Sidak post-test. *p<0.05, **p<0.01, ***p<0.001. ****p<0.0001.

**Supplemental Figure 6**

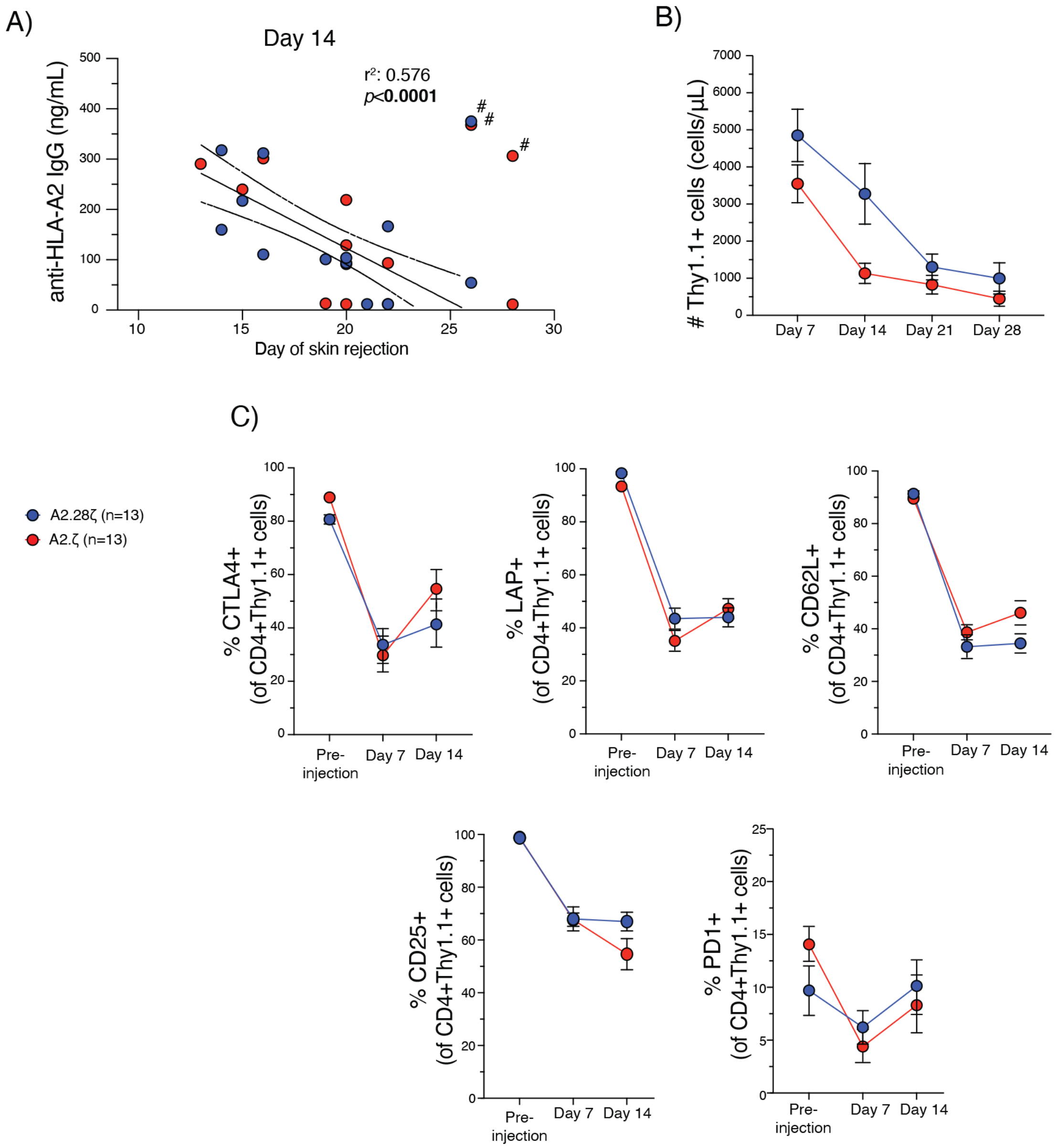


**Supplementary Figure 6:** *Supplementary material for the in vivo comparison of first- and second-generation CARs*. BL/6 mice were transplanted with skin grafts from syngeneic or HLA-A2$^{+}$ BL/6 mice and were intravenously administered 1x10$^{6}$ CAR-Tregs. (**A**) Levels of anti-HLA-A2 IgG Abs from mice infused with Tregs expressing first- and second-generation CARs. Data from PBS is shown. # Outliers values excluded from linear regression analysis. (**B**) Counts of Thy1.1$^{+}$ CAR-Tregs of total CD45$^{+}$ T cells in peripheral blood over time. (**C**) Phenotype of Thy1.1$^{+}$CD4$^{+}$ CAR Tregs in peripheral blood over time including expression of CTLA-4, LAP, CD62L, CD25 and PD1. Data are mean±SEM. Data pooled from 3 individual in vivo experiments. Statistical significance was determined using Pearson Correlation (**A**) and two-way ANOVA with a Holm-Sidak post-test (**B,C**). *p<0.05, **p<0.01, ***p<0.001. ****p<0.0001.

**Supplemental Figure 7**

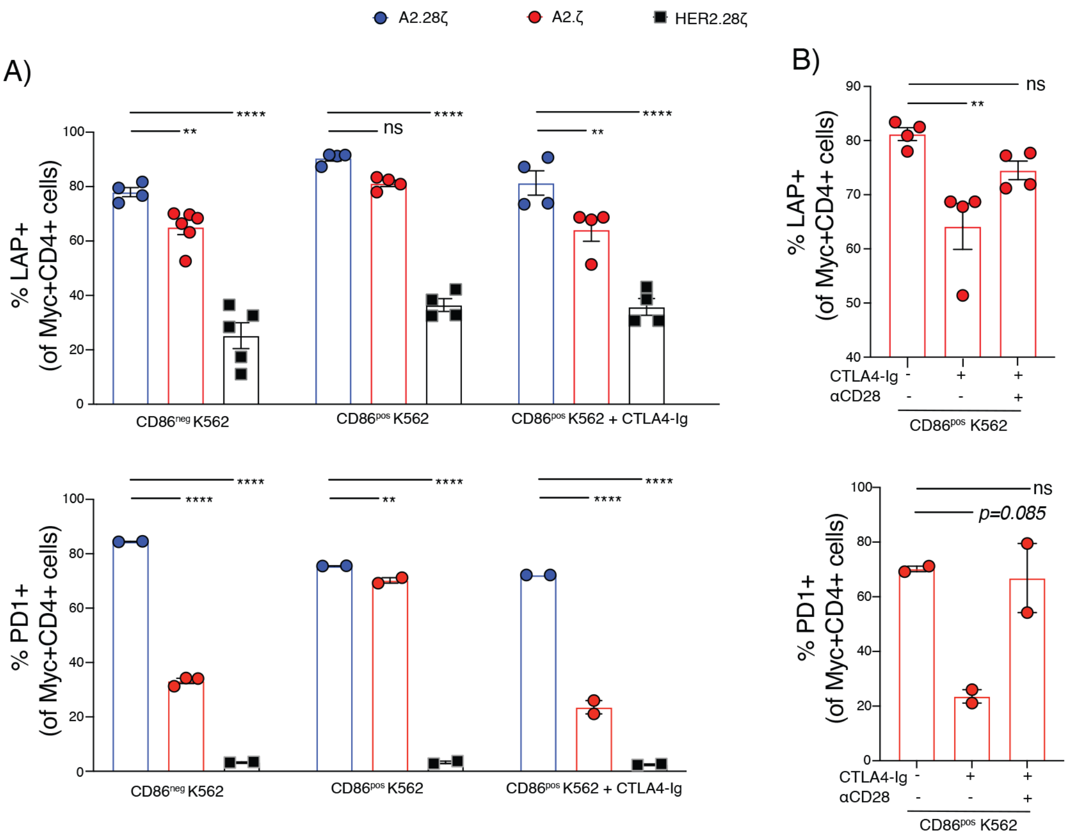


**Supplementary Figure 7:** *Supplementary material of the effects of a CAR-independent stimulation over the CAR-mediated activation and function of Tregs.* CAR-Tregs were co-cultured with $CD86^{+}HLA\text{-}A2^{+}$ or $CD86^{-}HLA\text{-}A2^{+}$ K562 cells at a 1:2 K562:Tregs ratio for 3 days. (**A-B**) LAP and PD1 expression in CAR-Tregs following 3-days of co-culture, gated on Myc+CD4+ live cells. Where indicated, CTLA4-Ig or activating anti-CD28 mAbs were added at 10 µg/mL. Data pooled from 2 or 1 independent experiments for LAP and PD1, respectively Data is shown as mean±SEM. Statistical significance was determined using one-way (**B**) or two-way (**A**) ANOVA with a Holm-Sidak post-test, *p<0.05, **p<0.01, ***p<0.001, ****p<0.0001.

**Supplemental Figure 8**

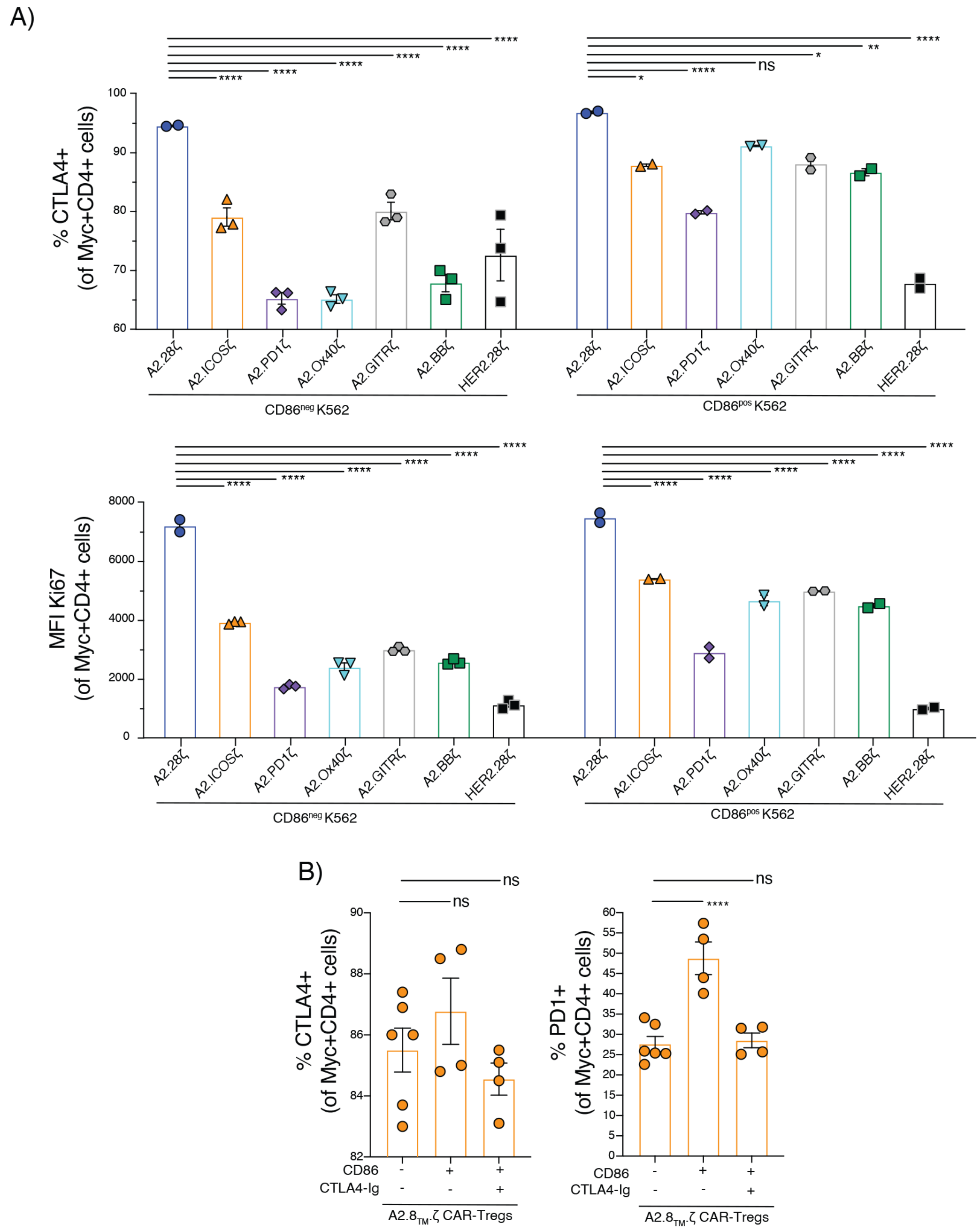


**Supplementary Figure 8:** *Supplementary material of the effects of a CAR-independent stimulation over the CAR-mediated activation and function of Tregs.* CAR-Tregs were co-cultured with CD86$^{+}$HLA-A2$^{+}$ or CD86$^{-}$HLA-A2$^{+}$ K562 cells at a 1:2 K562:Tregs ratio for 3 days. (**A**) CTLA4 and KIi67 expression in different co-stimulatory CAR-Tregs following 3-days of co-culture, gated on Myc+CD4+ live cells. (**B**) CTLA4 and PD1 expression in CD8α-TM CAR-Tregs following 3-days of co-culture, gated on Myc+CD4+ live cells. Where indicated, CTLA4-Ig or activating anti-CD28 mAbs were added at 10 μg/mL. Data pooled from 1 independent experiment. Data are shown as mean±SEM. Statistical significance was determined using one-way (**B**) or two-way (**A**) ANOVA with a Holm-Sidak post-test, *p<0.05, **p<0.01, ***p<0.001, ****p<0.0001.

**Supplemental Figure 9**

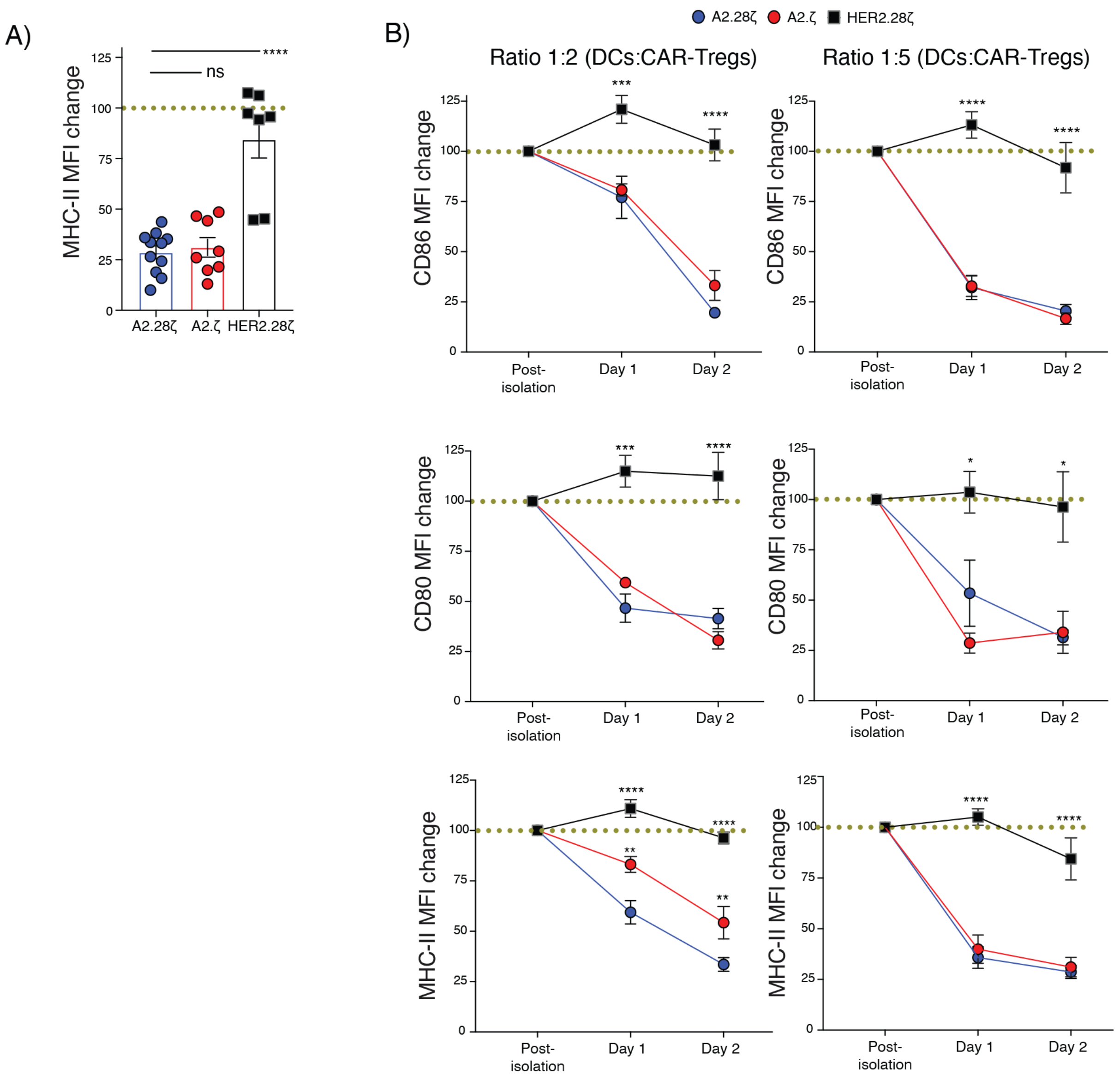


**Supplementary Figure 9:** *Supplementary material of the APC suppression by first- and second-generation CAR-Tregs.* CAR-Tregs were co-cultured with splenic HLA-A2$^+$ CD11c$^+$ DCs at a 1:2 or 1:5 DCs:Tregs for 1 or 2 days. (**A**) Expression of MHC-II in HLA-A2+ CD11c$^+$ DCs either immediately post isolation or cultured for 2-days with the indicated CAR-Tregs. (**B**) Expression of CD80, CD86 and MHC-II in HLA-A2+ CD11c$^+$ DCs cultured at different time points and with different ratios of DCs/CAR-Tregs. Data pooled from at least 4 individual experiments. Data are shown relative to DCs cultured with untransduced Tregs which were normalized to 100% (dotted lines). Data are shown as mean±SEM. Statistical significance was determined using one-way (**A**) or two-way (**B**) ANOVA with a Holm-Sidak post-test, *$p<0.05$, **$p<0.01$, ***$p<0.001$, ****$p<0.0001$.

**Supplementary Tables:**

**Supplementary Table 1**

| | **Group A** | **Group B** | **Pairwise significance (log-Rank)** |
|---|---|---|---|
| **CD28 family receptor CARs** | CD28 | ICOS CAR-Tregs | 0.427 |
| | CD28 | PD1 CAR-Tregs | 0.839 |
| **TNFR family receptor CARs** | CD28 | OX40 CAR-Tregs | 0.287 |
| | CD28 | 4-1BB CAR-Tregs | 0.202 |
| | CD28 | GITR CAR-Tregs | 0.543 |

**Supplementary Table 2**

| **CAR** | **Specificity** | **Hinge** | **Transmembrane domain** | **Costimulatory domain** |
|---|---|---|---|---|
| A2.28ζ | HLA-A2 | CD8α-derived | FWALVVVAGVLFCYGLLVTVA<br>LCVIWT | NSRRNRLLQSDYMNMTPRRPGLTR<br>KPYQPYAPARDFAAYRP |
| A2.ICOSζ | HLA-A2 | CD8α-derived | PVGCAAFVVVLLFGCILIIWF | SKKKYGSSVHDPNSEYMFMAAVNT<br>NKKSRLAGVTS |
| A2.PD1ζ | HLA-A2 | CD8α-derived | FWALVVVAGVLFCYGLLVTVA<br>LCVIWT | AVFCSTSMSEARGAGSKDDTLKEEP<br>SAAPVPSVAYEELDFQGREKTPELP<br>TACVHTEYATIVFTEGLGASAMGRR<br>GSADGLQGPRPPRHEDGHCSWPL |
| A2.CTLA4ζ | HLA-A2 | CD8α-derived | FWALVVVAGVLFCYGLLVTVA<br>LCVIWT | AVSLSKMLKKRSPLTTGVYVKMPP<br>TEPECEKQFQPYFIPIN |
| A2.GITRζ | HLA-A2 | CD8α-derived | LTVIFLVMAACIFFLTTVQLG | LHIWQLRRQHMCPRETQPFAEVQLS<br>AEDACSFQFPEEERGEQTEEKCHLG<br>GRWP |
| A2.OX40ζ | HLA-A2 | CD8α-derived | AFAVLLGLGLGLLAPLTVLLAL<br>YLL | RKAWRLPNTPKPCWGNSFRTPIQEE<br>HTDAHFTLAKI |
| A2.BBζ | HLA-A2 | CD8α-derived | TLFLALTSALLLALIFITLLF | SVLKWIRKKFPHIFKQPFKKTTGAA<br>QEEDACSCRCPQEEEGGGGGYEL |
| A2.TNFRζ | HLA-A2 | CD8α-derived | ISLPIGLIVGVTSLGLLMLGLVN<br>CIILVQR | KKKPSCLQRDAKVPHVPDEKSQDA<br>VGLEQQHLLTTAPSSSSSSLESSASA<br>GDRRAPPGGHPQARVMAEAQGFQE<br>ARASSRISDSSHGSHGTHVNVTCIVN<br>VCSSSDHSSQCSSQASATVGDPDAK<br>PSASPKDEQVPFSQEECPSQSPCETT<br>ETLQSHEKPLPLGVPDMGMKPSQA<br>GWFDQIAVKVA |
| A2.ζ | HLA-A2 | CD8α-derived | FWALVVVAGVLFCYGLLVTVA<br>LCVIWT | |
| HER2.28ζ | HER2 | CD8α-derived | FWALVVVAGVLFCYGLLVTVA<br>LCVIWT | NSRRNRLLQSDYMNMTPRRPGLTR<br>KPYQPYAPARDFAAYRP |
| A2.8$_{TM}$.28ζ | HLA-A2 | CD8α-derived | IWAPLAGICVALLLSLIITLI | NSRRNRLLQSDYMNMTPRRPGLTR<br>KPYQPYAPARDFAAYRP |
| A2.8$_{TM}$. ζ | HLA-A2 | CD8α-derived | IWAPLAGICVALLLSLIITLI | |
| HER2.8$_{TM}$.28ζ | HER2 | CD8α-derived | IWAPLAGICVALLLSLIITLI | NSRRNRLLQSDYMNMTPRRPGLTR<br>KPYQPYAPARDFAAYRP |

**Supplementary Table 3: List of antibodies used in the study**

| Target | Clone | Fluorophore | Company |
|---|---|---|---|
| CD4 | GK1.5 | BV785 | BD Biosciences |
| CD4 | RM4-5 | BUV563 | BD Biosciences |
| CD45 | 30-F11 | BUV805 | BD Biosciences |
| LAP | Tw7-16B4 | PerCP-e710 | BD Biosciences |
| PD1 | RMP1-30 | BV711 | BD Biosciences |
| CD25 | PC61 | BUV395 | BD Biosciences |
| CD80 | 16-10A1 | FITC or BUV395 | BD Biosciences |
| CD86 | GL1 | APC | BD Biosciences |
| CD8 | 53-6.7 | PE or AF647 | BD Biosciences |
| Thy1.1 | HIS51 | PE or BUV496 | eBioscience |
| Thy1.2 | 53 2.1 | BV510 or PECy7 | eBioscience |
| Ki67 | SolA15 | AF700 | eBioscience |
| FoxP3 | FJK-16s | FITC or PECy7 | eBioscience |
| CD11c | N418 | SB780 | eBioscience |
| Helios | 22F6 | PE-Dazzle | Biolegend |
| CTLA4 | UC10-4B9 | BV605 | Biolegend |
| I-A/I-E (MHC-II) | MS/114.15.2 | BV605 | Biolegend |
| CD62L | MEL-14 | AF647 | Biolegend |
| cMyc | 9E10 | AF647 or AF488 | UBC Ablab |
